# Unperturbed dormancy recording reveals stochastic awakening strategies in endocrine treated breast cancer cells

**DOI:** 10.1101/2021.04.21.440779

**Authors:** Dalia Rosano, Emre Sofyali, Heena Dhiman, Diana Ivanoiu, Neil Slaven, Sung Pil Hong, Andrea Rocca, Sara Bravaccini, Sara Ravaioli, Roberta Noberini, Tiziana Bonaldi, Luca Magnani

## Abstract

Hormone dependent breast cancer (HDBC) is the most commonly diagnosed tumor type in women. Adjuvant endocrine therapies (ET) have been the cornerstone in the clinical management of HDBC patients for over forty years. A vast proportion of HDBC patients incur long periods of clinical dormancy following ET, with tumour awakening appearing at a steady pace for up to 25 years (Pan et al., 2017). Extensive genomic studies have demonstrated that 15-30% of clinical relapses develop recurrent genomic changes which contribute to drug resistance (i.e. ESR1 activating mutations) (Bertucci et al., 2019; Magnani et al., 2017; Razavi et al., 2018). However, even in these cases, there is no conclusive evidence around the pre-existence vs. *de novo* nature of these events. We previously showed that ETs can trigger and select for dormancy in subpopulations of breast cancer (Hong et al., 2019). In this work we took two novel approaches to investigate the dormancy and awakening roadmap of HDBC cells at unprecedented detail. Firstly, we leveraged a rare cohort of n=5 patients which were treated with primary adjuvant ETs in the absence of surgery (TRACING-HT) to dissect the contribution of genomic aberrations to tumor awakening. Next, we developed a first of its kind evolutionary study *in vitro* to systematically annotate cancer cells adaptive strategies at single cell level in unperturbed systems during a period of several months (TRADITIOM). Collectively our data suggest that ETs steer HDBC cells into an inherently unstable dormant state. Over time, routes to awakening emerge sporadically and spontaneously in single lineages. Each dormant cell retains an intrinsic awakening probability which we propose is a function of epigenetic decay. Awakening occurs without an external trigger and involves multiple apparent endpoint phenotypes that cannot be fully explained by conventional Darwinian genetic selection processes. Finally, our data show that common genetic hits associated with resistance happen downstream of awakening. Overall, our data have uncovered previously unsuspected roles for stochastic nongenetic events during dormancy with profound clinical implications.

## Main

Understanding cancer cell dormancy remains one of the most enigmatic and fundamental challenges in cancer research as it is considered a driver of metastatic relapse, therapy resistance and immune evasion (Phan and Croucher, 2020). The contribution of tumour dormancy to disease progression is particularly evident in hormone dependent breast cancer (HDBC). All HDBC patients receive adjuvant endocrine therapies (ETs), which leads to clinical disease management, yet in many patients, cancer cells survive in a dormant state for years to eventually facilitate relapse and metastatic outgrowth. We reasoned that ET might induce cell cycle arrest and dormancy in residual micro-disseminated cancer cells (Hong et al., 2019). Mutational processes are by definition minimized in non-cycling cells, given the absence of DNA replication. This might possibly circumvent classical Darwinian evolution, which relies on randomly generated genetic variation and non-random selection. Therefore, we hypothesize that exit from dormancy (awakening) might have non-genetic factors. To gain an unprecedented look at this process, we first applied a high coverage WGS strategy (±100X) to a patient with bilateral metastatic HDBC managed by long-term primary ET (AI, Fig. 1A). This patient had extensive phenotypic heterogeneity at presentation (Supplementary Fig.1). Radiological examination shows that both lesions initially responded to therapy (6 months), followed by ±12 months of stable disease (putative dormancy) before both tumours progressed (awakening) whilst under therapy (48 months) (Supplementary Fig. 2). At this stage the patient accepted surgery and the four samples were profiled with WGS. In our work we define events which predate treatment as pre-existent, and events which occur during treatment as *de novo.* Focusing on pathogenic single-nucleotide variations (SNVs) in *bona fide* breast cancer drivers (FATHMM >0.6, (Martínez-Jiménez et al., 2020; Shihab et al., 2015)) could not identify high confidence SNVs compatible with progression (Fig. 1B). 36 months after surgery, the patient experienced a loco-regional relapse characterized by mutations in five *de novo* potential drivers, including ESR1 D538G (Razavi et al., 2018) and an additional FGFR2 V565L potentially explaining progression to the AI-FGFRi combination therapy. Next, we analysed an additional n=4 cases with very similar clinical histories (long-term management by AI in the absence of surgery) (Fig. 1C). In agreement with Patient A, we could not identify *de novo* breast cancer drivers at the time of tumour awakening (Fig. 1C). Extending the analysis to all known cancer drivers with less stringency on mutational filters does not result in additional candidates (Supplementary Fig. 3). Results from this rare cohort support our initial hypothesis that HDBC awakening might have a non-genetic component.

**Figure 1.**
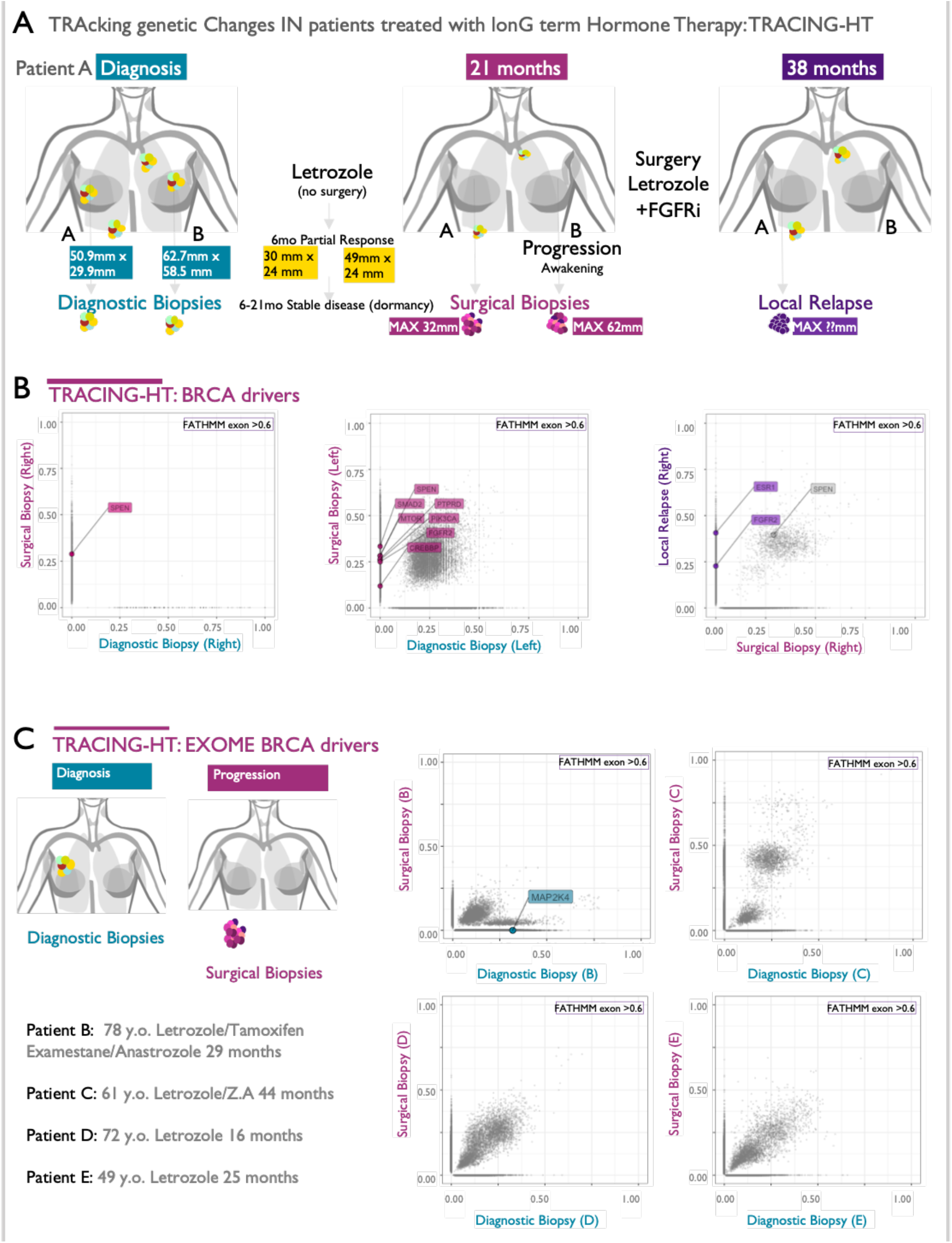
Whole genome profile of tumor awakening in the clinical setting. **A.** Clinical history of Patient A. **B.** Variant Allele Frequency plots from Patient A WGS data. Pairwise comparisons were done for pre-treatment and post-progression left and right lesions and post-progression vs. relapse for the right lesion. **C.** Variant Allele Frequencies plots from additional four patients’ WGS data. All patients were managed with primary endocrine therapy in the absence of surgery. Labelled genes passed two filters: bona fide breast cancer driver and FATHMM significant score >0.6 (predicted damaging). Variants detected in driver genes are labelled and highlighted according to detection in surgical biopsy (pink), diagnostic biopsy (cyan) or both (grey).

To quantify the role of non-genetic adaptive processes operating during ET-induced tumor dormancy we developed a trackable long-term *in vitro* study (**TR**acking **A**daptation, **D**ormancy and awaken**i**ng with mul**ti**-**OM**ics, TRADITIOM) (Fig. 2A).

**Figure 2.**
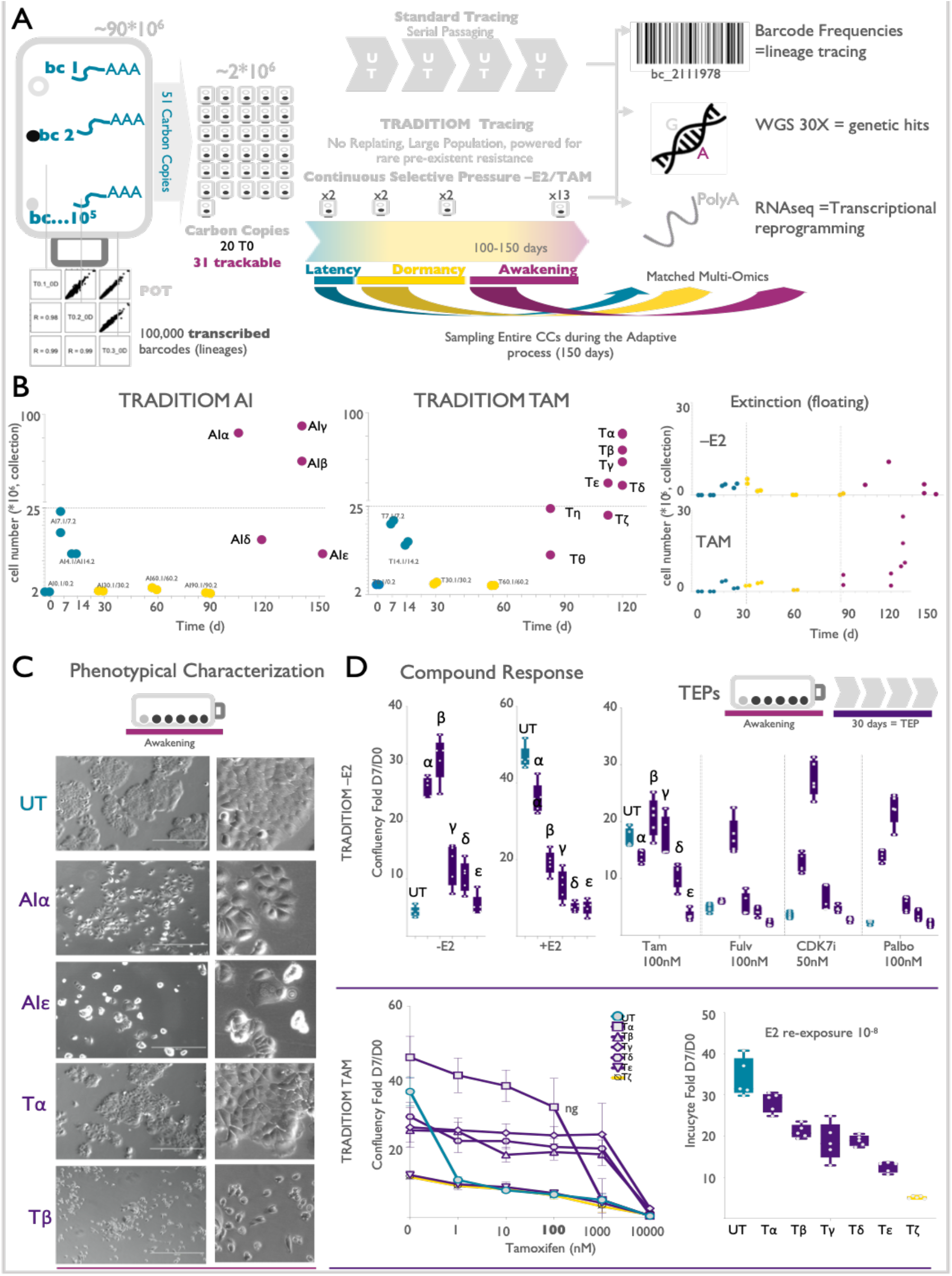
Longitudinal tracing via transcriptional barcodes. **A.** Schematic representation of Tracking adaptation, dormancy and awakening with multi-Omics (TRADITIOM) experimental setup. 1*10^5^ differentially barcoded cells were expanded to a founder population of 90*10^6^. Cells were replated in 31 trackable HYPERflasks (carbon copies), exposed to either –E2 (AI) or TAM and never passaged during the whole experimental course. In parallel, cells were kept in culture as untreated (UT) counterpart and were subjected to serial passaging. Part of the founder population was collected in 3 POT samples to analyse the initial barcode composition and replated in 20 time zero (T0) samples (reflecting the seeding of HYPERflasks) to determine the initial overlap between barcode composition in carbon copies. Harvesting of each HYPERflask was performed at the indicated time points (shared timepoints for 2 replicates for latency and dormancy and diverging time points for individual awakenings). Collected samples were analysed to reconstruct lineage adaptation dynamics (genomic barcode analysis), genetic contribution to therapy resistance (WGS) and transcriptional reprogramming under adaptation (RNA-seq) **B.** Cell counts in HYPERflasks for AI and TAM arm at their respective time of collection (teal: latency, yellow: dormancy, violet: awakening) were determined for both attached and floating cells. **C.** Representative bright-field images depicting morphological differences between untreated (UT) and awakening samples were captured with EVOS Cell Imaging Systems (10X). **D.** Growth rates of AI TEPs in response to oestrogen deprivation (-E2), re-exposure to oestradiol (E2) or treatment with different drugs: Tamoxifen (Tam, 4-OHT), Fulvestrant (Fulv), CDK7 inhibitor (CDK7i), Palbociclib (Palbo). Bottom panel: tamoxifen resistance analysis of the TAM TEPs to increasing doses of 4-OHT and re-exposure to E2. Representative graphs are shown as normalized confluency fold change upon 7 days of compound treatment (n=3).

TRADITIOM tracked MCF7 HDBC cells tagged with expressed genetic barcodes (100,000 poly-A barcodes) for 6 months in large Hyper-Flasks in the absence of cell passaging (Acar et al., 2020) (Fig. 2A and Supplementary Fig. 4A). TRADITIOM design minimizes all the confounding factors associated with lineage tracing studies (i.e. cell passaging and short term follow up (Hinohara et al., 2018; Zhang et al., 2021) and increase resolution down to single cell level to capture granular events associated with state transitions (cycling to dormancy and dormancy to awakening) (Acar et al., 2020). At a fundamental level, TRADITIOM allows us to map with unprecedented confidence all cell intrinsic awakening events in large population of cells. Ultra-low MOI (MOI=0.01) allows for quantitative estimation of heterogeneous cell cycle dynamics potentially occurring in cell lines (Supplementary Fig. 5). The founder population (90×10^6^ cells, POT) was sub-sampled in 31 identical trackable carbon-copies(~2×10^6^), to allow for parallel evolution leading to consecutive endpoints in response to different long-term treatments (Fig. 2A and Supplementary Fig. 4). Carbon-copies were then subjected to either oestrogen deprivation (AI) or Tamoxifen treatment (TAM), mimicking the standard of care for the current adjuvant ETs setting. An additional arm of the study followed untreated (UT) barcoded MCF7 during the same period of time with the usual cell passaging to examine changes occurring under neutral drift. Barcode analysis on three subsampled POT replicates and 20 time-zero (T0) carbon copies shows that plate parallelization introduces minimal changes in the starting barcoded cell population (Supplementary Fig. 4B). These data therefore demonstrate that the starting conditions of all carbon-copies are highly comparable. All the carbon-copies treated with ET exhibit an initial growth phase (latency period) followed by a rapid decrease in total cell number, which is confirmed by an increased number of floating dead cells (Fig. 2B). Cell counting analysis shows that AI and TAM carbon-copies reach a state comparable to dormant minimal residual disease (MRD) in about 30 days (Fig. 2B). Cells persist in a treatment induced dormancy state up to 150 days in the AI arm and up to 120 days in the TAM arm. Unexpectedly, the cell population in each flask begins to expand (awakening) in an asynchronous fashion, despite the almost identical starting conditions (Fig. 2B, Supplementary Fig. 6A). These observations are particularly robust in the AI arm (compare AIα to AIε). These data strongly suggest that the adaptive journey of each flask diverges with time. These findings are further corroborated by the radical morphological differences identified in awakened cells (Fig. 2C and Supplementary Fig. 6B). In order to further assess the adaptation trajectories of awakened samples, we passaged cells in the presence of ETs for an additional 30 days (Terminal End Points=TEPs) (Supplementary Fig. 6C). Tamoxifen resistance analysis of the TAM arm TEPs shows that population tolerance to increasing doses of 4-OHT (4-hydroxytamoxifen, Tam) varies among different aliases (Fig. 2D, bottom panels). Similarly, the AI arm shows different growth rates under oestrogen deprivation (Fig. 2D, upper panels). AI TEP samples also demonstrate distinct responses to most second-line treatment drugs such as Fulvestrant, Palbociclib (CDK4/6 inhibitor) and CDK7 inhibitor in combination with oestrogen deprivation (Fig. 2D). Re-exposure to E2 elicits discordant responses in both treatment arms. These data suggest that the phenotypes associated with awakening strongly diverge along the dormancy roadmap (Fig. 2D).

The asynchronous awakening observed in TRADITIOM is compatible with *de novo* events contributing to the switch from dormancy to cycling. Awakening carbon copies have approximately 3500, 4000 and 15000 surviving barcodes (untreated, -E2 and TAM, respectively) with the lower number of barcodes in untreated condition reflecting stochastic loss by serial passaging (Fig. 3A and Supplementary Fig. 7A). Around 40% of surviving barcodes were found to be in common between untreated carbon copies after 5 months (Fig. 3B). Similar overlap is observed in the AI arm, while TAM has marginally higher overlaps (Fig. 3B). These data show awakening is not mediated by a rare set of resistant cells, but many different lineages can potentially contribute to tumor progression over the course of months.

**Figure 3.**
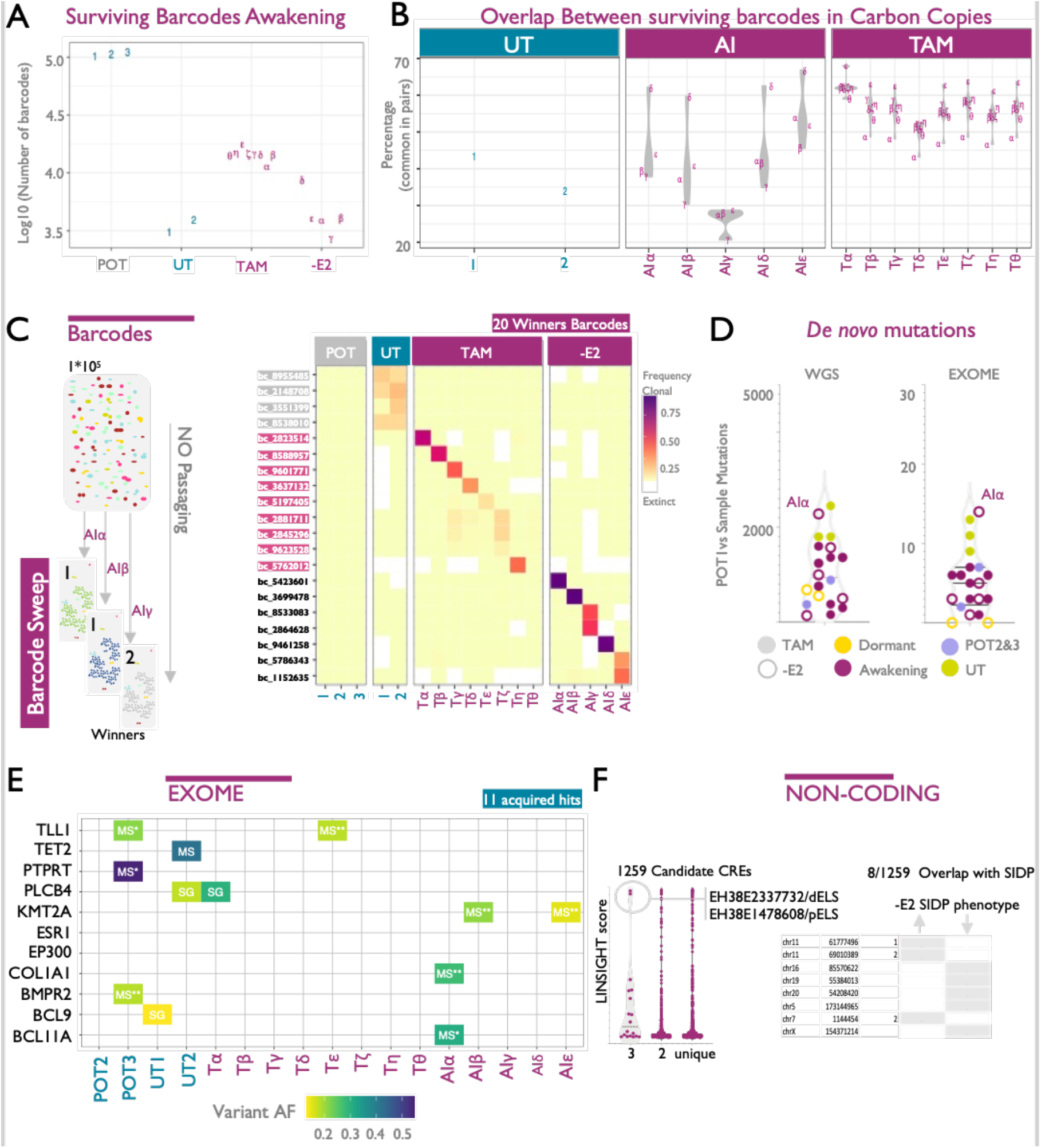
Awakening is a stochastic nongenetic process. **A.** Total number of barcodes in founder POT population and surviving barcodes in UT, TAM and AI (-E2) arm at the time of awakening. **B.** Percentage of common barcodes between two replicates for UT and awakening TAM and AI (-E2) samples. **C.** Heatmap of barcodes with highest frequencies (winners) among UT samples and TRADITIOM carbon-copies for both TAM and –E2 arm at the time of awakening. **D.** Number of total (WGS) and coding (Exome) *de novo* mutations compared to POT1 as determined by whole genome sequencing (30X) in dormant and awakening samples of TRADITIOM TAM and –E2 arm as well as UT counterparts. **E.** Acquired coding hits with their respective variant allele frequencies (FATHMM score >=0.1) (MS:missense, SG:stop-gain) (*damaging predicted by FATHMM, **damaging predicted by both FATHMM and MetaSVM) **F.** Noncoding variants were selected via OpenCravat (all SNV excluding Exome). Noncoding SNVs were filtered with the ENCODE Cis Regulatory Element function and sorted for LINSIGHT score. Noncoding SNVs called in 3 carbon copies and with a LINSIGHT score >0.2 were overlapped with our unpublished noncoding CRISPR-KRAB Screen (Repression of CRE under oestrogen deprivation). Upper arrow means gRNA enriched under ET treatment while down arrow means gRNA was depleted. Numbers indicate gRNA guide scoring per region.

Barcode analysis of the untreated arm confirmed nonspecific drift, with a small set of barcodes slightly expanding their frequency in the endpoint collection (Fig. 3C and Supplementary Fig. 7B). The situation is significantly different in the -E2 and TAM carbon copies where only 1 or 2 awakening barcodes (winners) quickly sweep through the entire population (Fig. 3C). Conducting the experiment without additional perturbation therefore offers a radically different view from previous efforts (Hinohara et al., 2018), since our data allows us to capture unconventional evolutionary events beyond classical Darwinian competition between cycling clones (fitness advantages) but captures the awakening event and the lineage where it occurred. Our data show that 12 out of 13 carbon copies have winning barcodes, further demonstrating that awakening is not driven by pre-existent traits. In 2/13 carbon-copies (AIγ and AIε), two barcodes appear to awaken simultaneously. Extensive analysis suggests that this was not due to cells containing two barcodes and might indicate cooperative behaviour. TAM carbon copies have evidence of multiple lineage expansion (T**ζ** and Tγ) and one sample does not have clear winners (Tθ). Interestingly, Tθ cell count hinted the carbon copy had just emerged from dormancy (Fig. 2B). One limitation of our study is that the extensive barcode complexity (which was chosen to capture ultra-rare pre-existent lineages) allows for multiple individual barcodes to tag the same initial phenotypes. This would predict the existence of a common genetic ancestor carrying a specific genetic driver in all winning barcodes. We tested this hypothesis by matched WGS profiling of all the awakening samples. With the exception of AIα, the mutational burden of all ET flasks was lower than UT counterparts (Fig. 3D). These data therefore are in agreement with the hypothesis that dormancy decelerates mutational processes. In agreement with barcode analysis, carbon copies possessed unique sets of *de novo* coding hits (Fig. 3E). Examining all *bona fide* drivers shows that the UT arm appears to have equal chance of developing *de novo* genetic hits (Fig. 3E), while several samples have no evidence *de novo* coding events. These *de novo* events might also reflect the passive expansion of a pre-existent rare clone, as shown by calls in POT2, which likely reflects genetic heterogeneity in subsampled analysis. Importantly, none of the detected *de novo* variants have been described as drivers *in BRCA* (Martínez-Jiménez et al., 2020). Collectively, these data demonstrate that awakening events are not mediated by a *de novo* coding driver. Next, we investigated the potential role of *de novo* noncoding variants in the awakening process. We reasoned those likely candidates should be recurrent within our datasets. Restricting our analysis on ENCODE annotated regions we found 1252 candidates, shared between 1 to 3 carbon-copies (Fig. 3F). Only six CIS elements shared by at least 3 plates have a functional prediction score (LINSIGHT score) > 0.2. Furthermore, cross analysis with our SIDV platform (dCas9-KRAB perturbation of clonal MCF7 enhancers, unpublished) shows that these regions have likely no functional impact in MCF7. Overall, these data demonstrate that awakening is not mediated by a preexistent or *de novo* genetic hit.

TEPs provide invaluable insights into the long-term outcomes of the adaptive journey. All winner barcodes in the AI arm are preserved and consolidated in their TEP counterparts whereas the TAM arm reveals a more dynamic barcode evolution. Four of the awakening TAM carbon copies show winning barcode consolidating even further (α-β-ε-**ζ**), while two samples show significant sweeps between awakening and TEP (Tγ and T-δ, Fig. 4A-B).

**Figure 4:**
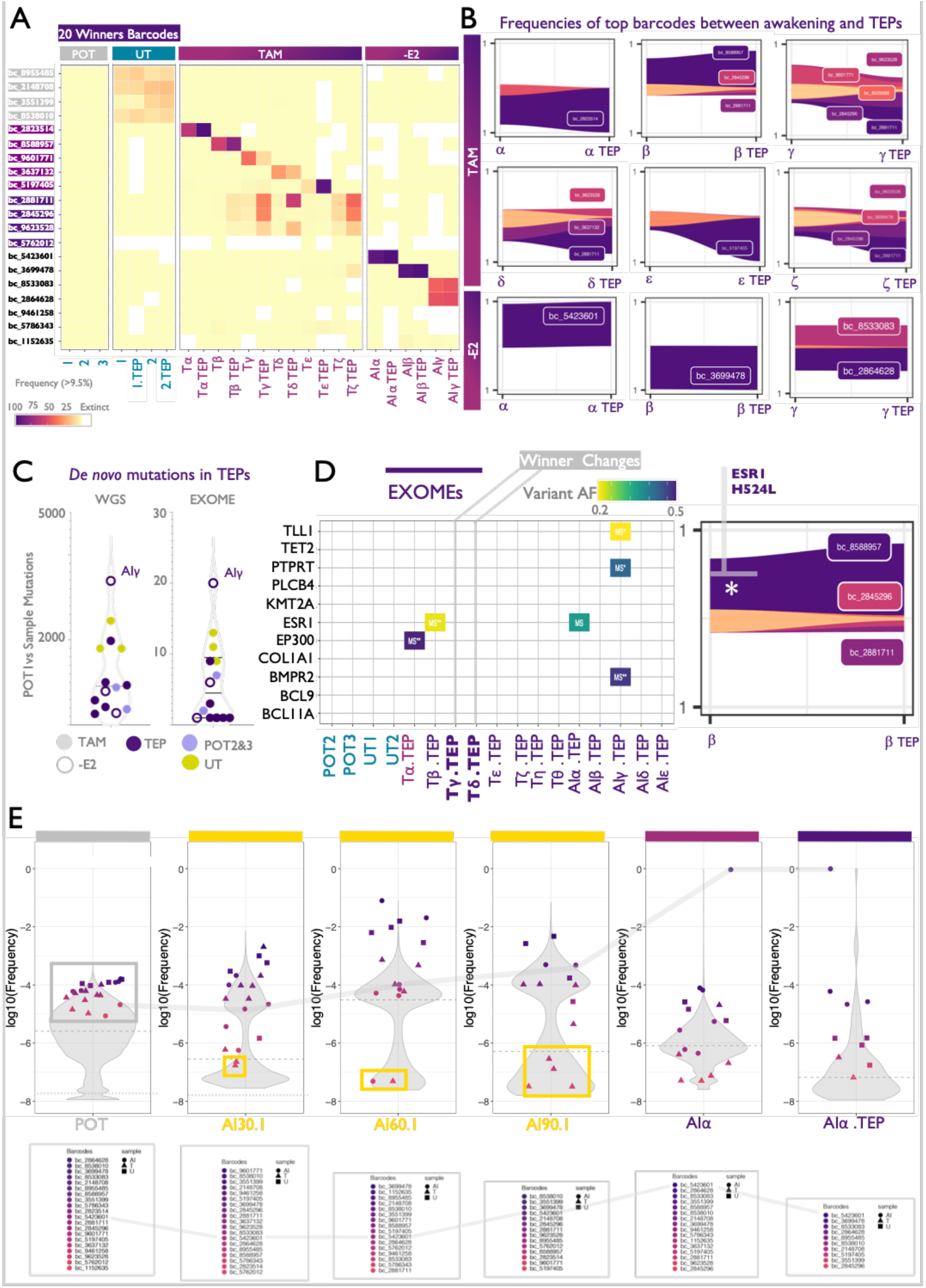
Awakening is a sequential process and it is not the final step of adaptation. **A.** Barcode enrichment analysis: Heatmap of barcodes with highest frequencies (winners) among UT samples and TRADITIOM carbon-copies for both TAM and –E2 arm at the time of awakening and their corresponding TEPs. **B.** Fish plots depicting the changes in frequency of winning barcodes in the journey between awakening and TEP of representative TAM and –E2 carbon copies. **C.** Number of total (WGS) and coding (Exome) *de novo* mutations compared to POT1 as determined by whole genome sequencing (30X) in TEP samples of TRADITIOM TAM and –E2 arm as well as UT counterparts. **D.** Acquired coding hits with their respective variant allele frequencies along with a depiction of *de novo* ESR1 mutation in TAMb TEP (FATHMM score >=0.1) (MS:missense, SG:stop-gain) (*damaging predicted by FATHMM, **damaging predicted by both FATHMM and MetaSVM). **E.** Violin plots depicting the frequency distribution of barcodes during their adaptive journey through POT, dormancy, awakening and TEP with a focus on bc_5423601, the winning barcode for AIα carbon copy (trajectory over time highlighted in grey). Winning barcodes (Panel A), at each specific stage, are indicated under the corresponding violin plot in order of frequency. Specific symbols (square for UT, circle for -E2 and triangle for Tam) indicate the belonging of the winning barcodes to the respective treatment.

Post-awakening carbon copies might reactivate Darwinian clonal competition between awakened cycling lineages. To test if the sweep between awakening and TEP is driven by *de novo* coding mutations, we again resorted to WGS but could not identify any significant changes between awakening and TEPs (Fig. 4C-D). Conversely, the Tβ carbon copy offered a glimpse of the temporal order connecting nongenetic and genetic events. In this particular case, cells acquire a non-canonical ESR1 H524L mutation, which has been previously shown to promote Tamoxifen resistance (Chung et al., 2017; Toy et al., 2019) after nongenetic awakening (Fig. 4C-D). While all AI awakening carbon copies have private winning barcodes, the TAM β, γ, δ and **ζ** TEPs show some evidence of common winner barcodes (Fig. 4A-B). WGS analysis rules out a common genetic driver shared among these carbon copies (Fig. 4D). These data suggest that AI awakening is purely stochastic while TAM awakening might combine stochastic awakening with pre-existent nongenetic selection.

To investigate longitudinal evolution of lineage composition, we explored the barcode frequency dynamics for all winners (top barcodes in all UT, -E2, TAM conditions) across the entire TRADITIOM study. Unexpectedly, all winning barcodes were found at high frequencies in POT (Fig. 4E and Supplementary Fig. 8). Indeed, the 4 barcodes expanding under neutral drift (UT arm) were found at the top of the distribution in most T0 carbon copies and successive timepoints (Riding the front of the wave, grey boxes, Supplementary Fig. 8 and 9). A similar trend is initially observed for TAM and -E2 specific winners where all winning barcodes are present at high frequencies in POT and T0. These data suggest that the initial barcode frequency might influences the probability of entering in dormancy. Nevertheless, in the ET arms, contrarily to UT dynamics (riding the wave) the transient barcode frequency stops becoming a reliable predictor once dormancy is set (day 30, yellow boxes, Figure 4E and Supplementary Fig. 9). Once lineages have passed the dormancy bottleneck, given sufficient time they all have a chance of awakening. This observation strengthens the notion that stochasticity might indeed play a key role in awakening. Winner barcodes switch between awakening and TEP in 2 TAM carbon copies suggests that runner-up high frequency barcodes present at awakening can outgrow at a certain step along adaptation, outcompeting the actual winning barcode, creating an unstable phenotype. Furthermore, these data strongly suggest that sequential awakening events occur spontaneously with comparable probabilities in surviving dormant cells via a non-genetic event contributing to an overall unstable phenotype. These results have a profound implication for the concept of the metastatic cascade, which is generally thought as a process of late-stage disease driven by existing deposits beginning to metastasize at increasing frequencies. Our data would offer an alternative explanation, where the first signs of metastatic progression act as a whistleblower for additional sequential awakening events in residual dormant clones.

Phenotypic, genomic and lineage tracing (Fig. 2-4) show that awakening is a rare (1-2 winning barcodes) and stochastic event occurring sequentially in residual dormant HDBC cells. Awakening cells exhibit unexpected phenotypic heterogeneity which lead us to hypothesize the existence of stochastically accessed heritable awakening mechanisms becoming available during dormancy.

We began investigating this question via 51 RNA-seq, matching the WGS and barcode analysis (Fig. 2A and 5A). One hundred and seventy days of neutral drift does not significantly alter the RNA-seq profile of MCF7 cells, with UT replicates having almost indistinguishable RNA profiles. This suggest that previously reported divergence is probably mediated by long-term lab-to lab culturing differences rather than continuous divergence (Ben-David et al., 2018). These data also suggest that the substrate of stochastic processes (i.e., epigenome decay or transposon re-activation) is either muted or inconsequential in the absence of selective pressure. Previous and unpublished scRNA-seq from our group show that ET triggers and selects dormant cells in a relatively short period of time (Hong et al., 2019). TRADITIOM confirms the rapid transcriptional effect of ETs, with TAM acting more quickly than oestrogen deprivation (Fig. 5A). In agreement with previous data, dormancy appears to involve a robust and reproducible bottleneck, which can last relatively unaltered for months (Fig. 5A). Interestingly, RNA profiles suggest that -E2 latency is relatively more protracted but 14 days of treatment begin to impart RNA profiles which strongly resemble fully dormant cells (Fig. 5A). In agreement with phenotypic, genomic and barcode analyses (Fig. 2C-D and 3C-E), AI awakening carbon copies are exceedingly heterogenous with TEP carbon copies continuing to evolve past the awakening event. TAM awakening carbon copies show a lower degree of divergence compared to AI but confirm the continuous adaptation process occurring in TEP. Collectively, these data strongly support the notion that ET corners cells in dormancy but awakening propels dormant cells into an unpredictable phenotypic landscape.

**Figure 5:**
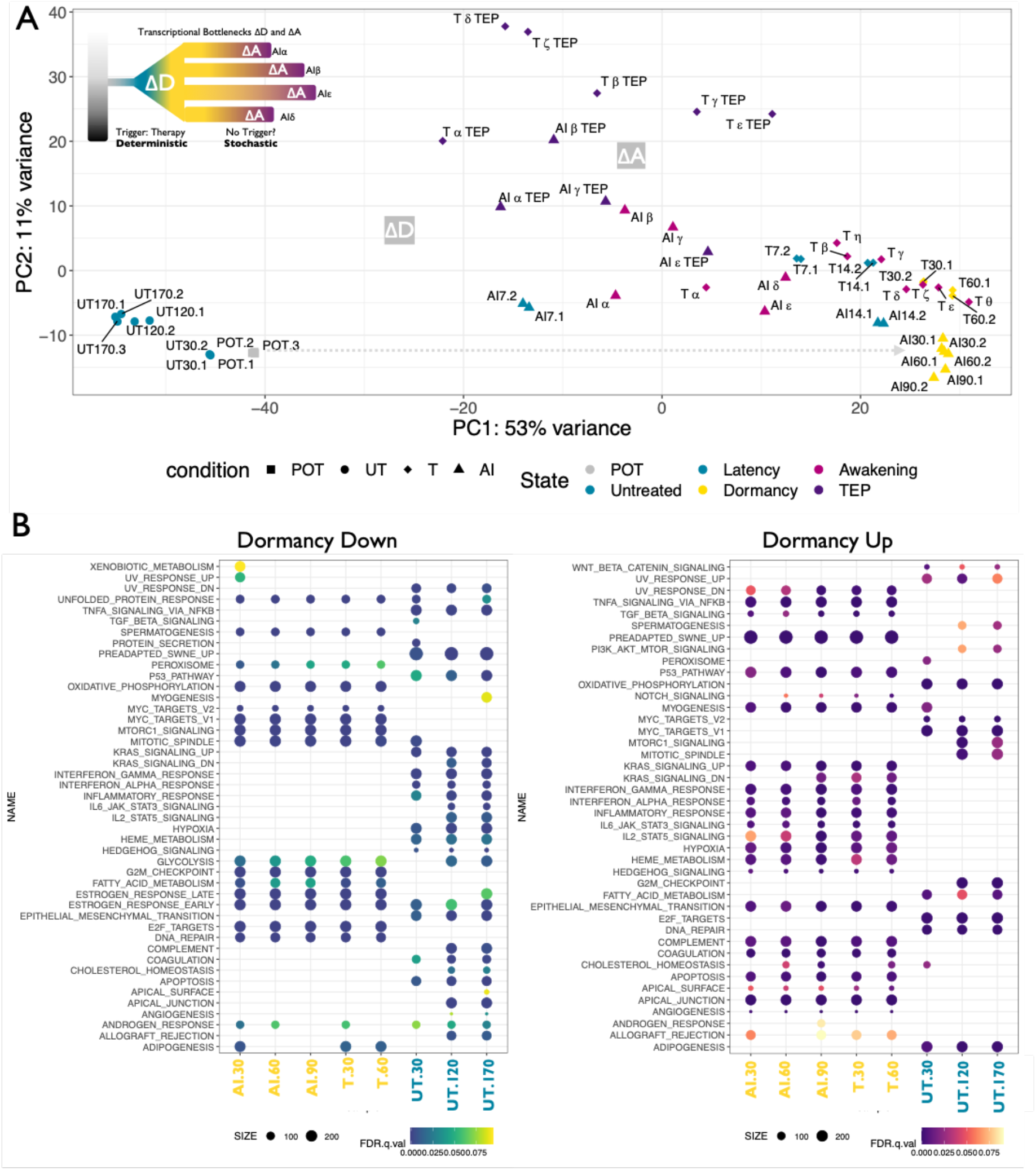
ET therapies trigger dormancy. **A.** Principal Component Analysis (PCA) of RNAseq-based expression data of all TRADITIOM dataset. **B.** Gene set enrichment analysis (GSEA) comparing POT to UT day 30, day120 and day170 and dormancy (AI day 30, 60, 90 and TAM day 30, 60) (FDR<=0.1). Enrichment of a gene set is depicted by a point colour coded based on FDR that supports the enrichment and size of the point corresponds to the number of genes observed as expressed in the data.

AI and TAM arms dormancy profiles (from day 30) were quite reproducible (Fig. 5A). During dormancy, the RNA-seq profile shows modest changes. Taken together with barcode analysis (Supplementary Fig. 8-9), these data suggest that cells do not gradually transition toward awakening but are truly awaiting for an awakening event. Gene Set Enrichment Analysis identifies clear pathways defining dormant cells (Fig. 5B). As expected, dormant cells suspend oestrogen and confirm the Pre-Adapted dormancy signature we have previously identified (Chen et al., 2021; Hong et al., 2019). As expected, dormant cells also suspend metabolic activities (glycolysis and oxidative phosphorylation), G2M checkpoint and MYC signalling, in agreement with quiescence (Rehman et al., 2021). Interestingly, while P53 activity is increased, DNA repair is downregulated, a situation compatible with lower mutational burden. On the other hand, dormant cells robustly activate pro-inflammatory pathways (NFKB, interferon response and IL6 (Hong et al., 2019; Sansone et al., 2016; Siersbæk et al., 2020) in addition to epithelial to mesenchymal transition and apical junction, suggesting complex hybrid states. Of note, a large scale CRISPR-dCas9 noncoding screen from our lab (unpublished) clearly identifies NFKB as a central regulator of dormancy entry. We then compared the common dormancy profile to the individual awakening RNA signatures (Fig. 6A).

**Figure 6:**
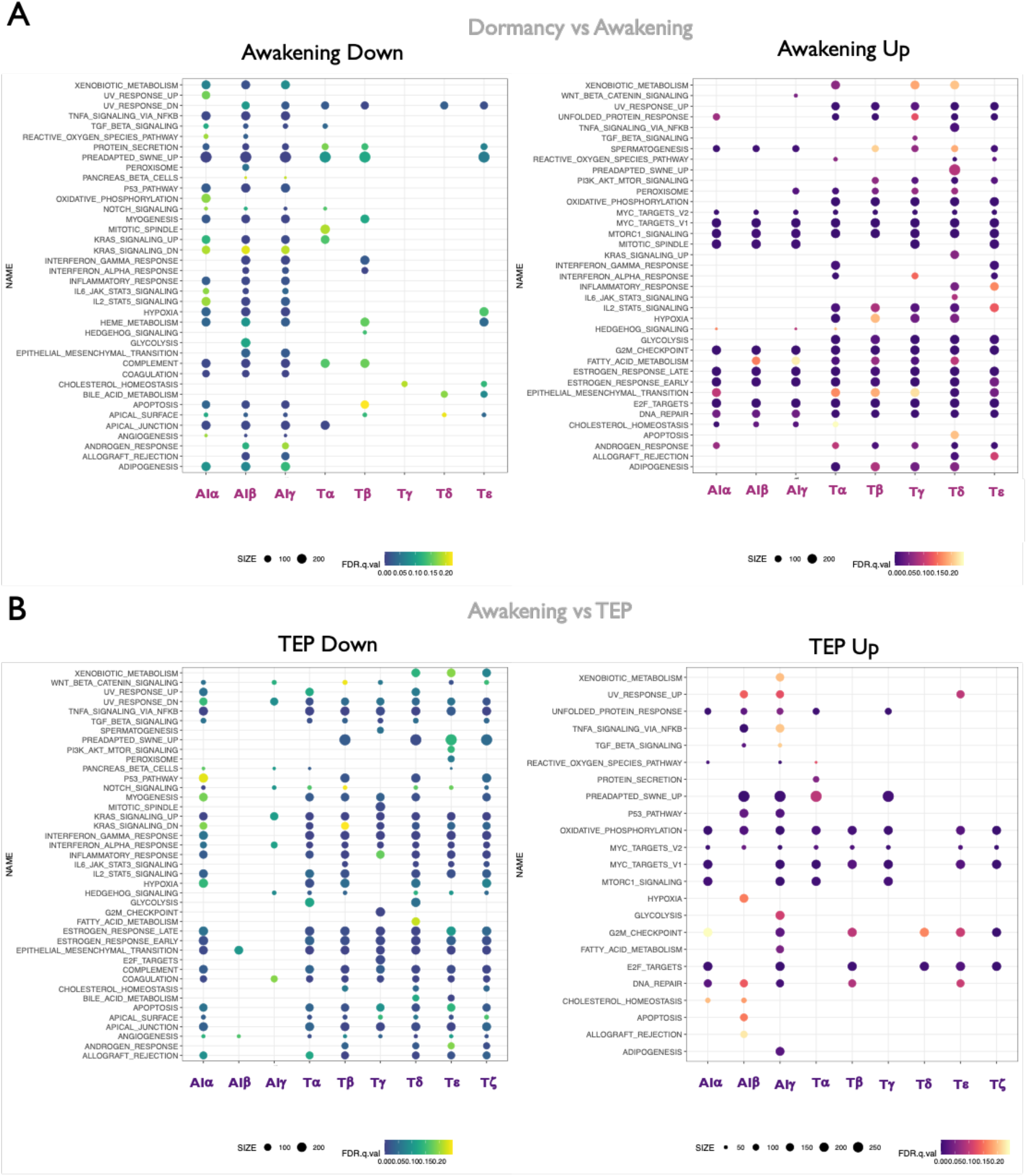
Awakening and TEP lead to unpredictable phenotypes. **A.** Gene set enrichment analysis (GSEA) comparing dormant to awakenings carbon copies (FDR<=0.25). **B.** Gene set enrichment analysis (GSEA) comparing awakenings to TEPs (FDR<=0.25). Enrichment of a gene set is depicted by a point colour coded based on FDR that supports the enrichment and size of the point corresponds to the number of genes observed as expressed in the data.

Many of the enrichment sets defining dormancy entrance and maintenance are completely reversed upon awakening, demonstrating extensive transcriptional reprogramming occurring in single lineages. However, the robust level of reproducibility common to dormant cells is not as obvious at awakening, confirming phenotypic divergence (i.e.peroxisome, Pre-adapted signature, Glycolysis). This divergence becomes more prevalent at TEP (Fig. 6B). These data suggest that even in relatively simple cell models, the phenotypes and the timing of awakening are highly unpredictable. Arguably, this suggests that developing a therapeutic strategy on the first awakening will not guarantee efficacy in additional awakening clones.

Finally, we took initial steps toward identification of the substrate which could connect time and stochastic awakening. In Darwinian clonal evolution, this is represented, at least in part, by stochastic genetic variation (Martincorena et al., 2018; Nik-Zainal et al., 2012, 2016; Petljak et al., 2019; Tomasetti and Vogelstein, 2015; Tomasetti et al., 2013, 2017) (i.e., Mutational Signature 1). We reasoned that in noncycling cells, the epigenome could be a source of stochastic change, for example passive DNA methylation forms the basis of the epigenetic (Horvath, 2013) clock. However, cell lines have lost most of their methylation due to culture condition (Liu et al., 2016; Roulois et al., 2015) so we focused our attention on epigenetic modifications, a well-established barrier to phenotypic divergence (Suva et al., 2013) with ties to ET resistance (Magnani et al., 2013; Nguyen et al., 2015). Leveraging unbiased mass spectrometry analysis, we show that dormant cells gain significant histone marks associated with heterochromatin state (i.e. H3K9me2/3, H3K27me3, H3K9me2/3 and H4K20me3) and lose most histone acetylation marks (i.e. H3K9acK14ac and H4Kpan ac) (Fig. 7A). For the most part, these changes are reversed in awakening cells (Fig. 7A) suggesting that epigenetic decay involving these modifications might be responsible for stochastic awakening.

**Figure 7:**
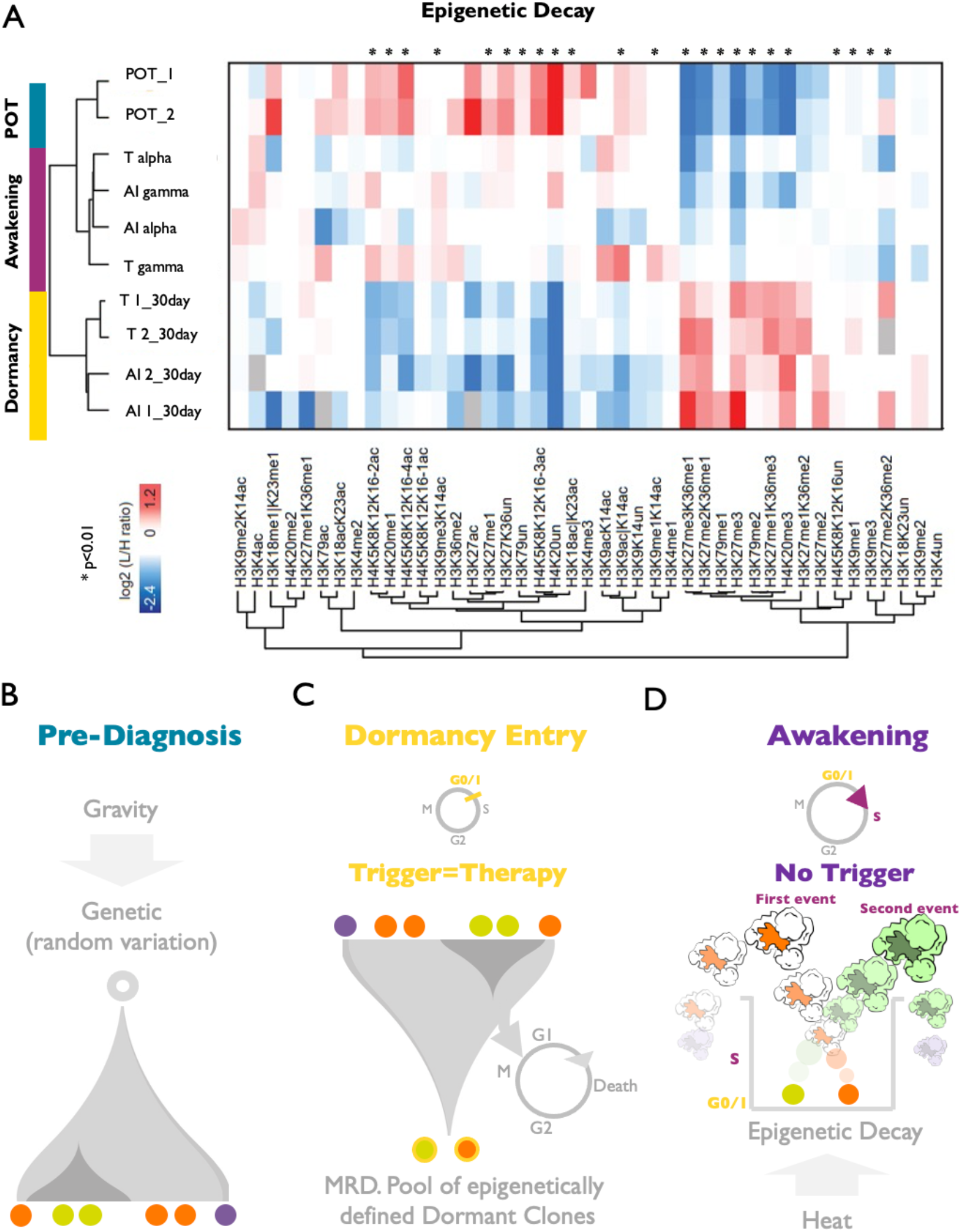
Model. **A.** Mass-spectrometry analysis of post-translational histone modifications in selected POT, dormancy and awakening samples. (*= p<0.01) **B.** Model Part I: Primary tumor heterogeneity is driven by natural selection of stochastic genetic variation. **C.** Model Part II: Adjuvant ET impose cell partly inheritable cycle restriction (dormancy) in actively proliferating cells. **D.** Model part III: the “dormancy pop-corn pot”. The “dormancy pot” limits accumulation of genetic mutations due to cell cycle arrest, but dormancy is epigenetically unstable (heat=epigenetic decay) and all dormant cells in the pot will sequentially “pop”. The first event, as the following ones, have limited clinical value since each awakening cell can be characterized by different phenotypes.

## Discussion

Distinct from almost all other cancer types, the clinical journey of patients with hormone dependent breast cancer (HDBC) is unequivocally shaped by tumor dormancy. HDBC tumors are clinically managed via adjuvant endocrine therapies (ETs), yet in many patients, cancer cells can survive in a dormant state for years before eventually causing clinical relapses and metastatic outgrowth. Our view is that ETs induces dormancy in most disseminated cancer cells. In up to ~50% of patients, dormancy extends long enough to transform biological dormancy into curative successes. However, in the remaining ~50% of patients, tumors eventually relapse with metastatic disease, at least in part as a consequence of tumor cell awakening from ET-induced dormancy. TRADITIOM is the first major conceptual study exclusively designed to chart the dormancy roadmap at single cell level in unperturbed systems (no cell replating) during endocrine therapy-induced dormancy and awakening. Results from TRADITIOM show that entrance into dormancy has deterministic triggers/phenotypes while exit from dormancy is an inevitable, rare, stochastic event. All the 31 carbon copies entered dormancy simultaneously after one month of exposure to ET (-E2 or TAM) with reproducible transcriptional profiles which remained relatively unchanged for months until awakening. Conversely, the stochastic awakening event (asynchronous awakening) is manifested via multiple dissimilar phenotypes that further diverge along the multistep roadmap toward resistance. Our data highlight the absence of a common genetic trigger for awakening, hinting that awakening leverages nongenetic processes (i.e. epigenetic decay) in the context of random genetic variation.

We propose a three-step model for cancer evolution in HDBC patients treated with ET (The pop-corn Model, Fig. 7B-D): in treatment naive women, exponentially growing cancer cells compete, and clonal evolution is fuelled by random genetic variation (unrepaired DNA damage and DNA polymerase errors=gravity) (Fig. 7B). This force is shaped by natural selection and results in tumor heterogeneity in untreated primary HDBCs. After diagnosis and surgery, endocrine therapies actively induce dormancy in surviving cells: the “dormancy pop-corn pot” (Fig. 7C). The “dormancy pot” limits genetic noise due to loss of cell cycling and lowered mutational processes. Dormancy is inherently unstable (epigenetic decays) and given infinite time, all dormant cells in the pot will sequentially “pop” (Fig. 7D) The first event has limited clinical value since each awakening cell can be characterized by alternative phenotypes. This model might explain clinical observations such as the “metastatic cascade” (as first awakening event (pop) predicts a closer second event and so on) and the conundrum of precision medicine in the relapse setting (multiple sequential phenotypes which cannot be counteracted). These findings have the potential to conceptually transform the therapeutic field as they highlight the utmost importance of developing actionable and scalable strategies to target cells entering dormancy, thus modifying their numbers and indirectly affecting awakening dynamics.

### Limitation of the study

Longitudinal lineage tracing analysis suggests a stochastic trajectory especially for awakening samples of AI treatment arm. On the other hand, the existence of common barcodes at awakening and corresponding TEPs of TAM arm does not exclude**_**the possibility of contributing pre-existent mechanisms at play. Our extensive whole genome sequencing analysis provides substantial evidence for the lack of a common genetic driver for TAM awakening. In order to dissect the individual contribution of transcriptional evolution and epigenetic inheritance to awakening at a single cell level, a miniaturized and modified version of TRADITIOM (TRADITIOM-LIGHT) with a lower barcode complexity and higher replication rate, is currently underway (four months running) where all lineages are being traced via scRNA-seq (transcribed barcodes) over the course of dormancy roadmap.

## Methods

### Cell culture

Breast cancer cell line MCF7 was kindly provided by Philippa Darbre (PMID: **26610607**). Cells were cultured in Dulbecco’s Modified Eagle Medium (DMEM) supplemented with 10% FCS, 50 units/ml penicillin, 50 μg/ml streptomycin sulfate (Invitrogen AG, Carlsbad, USA), and 10^-8^M 17-ß-estradiol (E2, Sigma-Aldrich, Saint-Louis, USA) and were kept at 37°C with 5% CO_2_ in a humidified atmosphere. Cells were routinely tested for Mycoplasma contamination.

### Cell culture in Hyperflasks

MCF7 cells were grown without passaging in High Yield PERformance Flasks (HYPERflask^®^) cell culture vessel (Corning, CLS10034) and maintained either in 100nM 4-Hydroxytamoxifen (TAM, Sigma-Aldrich H7904) or kept in phenol red-free DMEM (Gibco, 11880028) supplemented with 10% charcoal stripped FCS (-E2, estrogen deprivation). Medium was changed weekly and cells were monitored twice a week for apparent growth changes and harvested upon awakening (resumption of cell proliferation evaluated by visual inspection using EVOS cell imaging system, Thermofisher Scientific). Untreated arm of TRADITIOM was maintained in E2 supplemented media and underwent serial passaging (1:10 on average, twice a week).

### Floating cells harvesting

Floating (dead) cells were collected from media at each collection point from the large volume of media (550 ml) of HYPERflask culture system. MCF7 is an adherent cell line, with cells that detach from the flask surface upon death. By centrifuging the culture media at 1,200 rpm for 5 min in Corning 250 mL Centrifuge Tubes (Corning) we collected cells that had died within the week. Collected cells were counted with hemocytometer and trypan blue to measure cell viability.

### TRACING-HT

Extraction of DNA from fresh frozen (Buffy Coat and Drug Resistant Tumour) samples was carried out using DNeasy Blood and Tissue kit (Qiagen, 69506). Extraction from FFPE samples was conducted using GeneRead DNA FFPE Kit (Qiagen, 180134). Quality and quantity of DNA was determined using the TapeStation 2200 System (Agilent) with the Genomic DNA Screentape Analysis (5365). To improve the proportion of DNA fragments at the optimal length for library preparation samples were sonicated for 10 cycles using the Bioruptor Pico Sonication Device (Diagenode). FFPE DNA samples were treated with the NEBNext FFPE DNA Repair Mix (NEB, M6630L). DNA libraries for Illumina sequencing were prepared with the NEBNext Ultra 2 DNA Library Kit for Illumina (NEB, E7645L) using 200ng of DNA and custom made unique dual indices (8bp), a kind gift from Dr. Paolo Piazza (British Research Council Genomics Facility). DNA libraries were quantified using the TapeStation 2200 System with the High Sensitivity D1000 Screentape Analysis (Agilent, 5584). Samples were pooled based on type of the original material; FFPE, Fresh Frozen. Normal DNA was pooled at 1:3 ratio to tumour material. Pooled DNA libraries were sequenced withNovaSeq using the S2 50bp Paired End flow cell chemistry (Output 333-417 Gb).

Clinical informations Patient A:, 2018 Metastases – Fresh Frozen material. Patient B:, Normal - Buffy Coat – Fresh Frozen, Untreated Diagnostic Biopsy – FFPE, Drug Resistant Tumour – Fresh Frozen. Patient C:, Normal – Buffy Coat – Fresh Frozen, Untreated Diagnostic Biopsy – FFPE, Drug Resistant Tumour – Fresh Frozen. Patient D: Normal – Buffy Coat – Fresh Frozen, Untreated Diagnostic Biopsy – FFPE, Drug Resistant Tumour – FFPE. Patient E:, Normal – Buffy Coat – Fresh Frozen, Untreated Diagnostic Biopsy – FFPE, Drug Resistant Tumour – Fresh Frozen

### Cell barcoding

CloneTracker XP 10M Barcode-3’ Library with RFP-Puro (BCXP10M3RP-P) was purchased from Cellecta. Production of lentiviral particles and MCF7 transduction was performed following CloneTracker™ XP Lentiviral Expressed Barcode Libraries online manual (https://manuals.cellecta.com/clonetracker-xp-lentiviral-barcode-libraries/). Briefly, HEK-293T cells were transfected with cellecta CloneTracker XP library and ready-to-use lentiviral packaging plasmid mix (Cellecta, CPCP-K2A) using Lipofectamine (Thermo Fisher Scientific, 18324020) and Plus reagent (Thermo Fisher Scientific, 11514015). Viral particles were collected 48 hours upon transfection and precipitated overnight with PEG-IT Precipitation Solution (LV810A-1-SBI). Lentiviral titration was performed by flow cytometry. 10×10^6^ MCF7 cells were transduced with 0.01 MOI (multiplicity of infection) using 0.8 μg/ml polybrene to get a final number of 1×10^5^ differentially barcoded cells. For selection, 2.5μg/ml puromycin was added to the culture media for two cycles of 72 hours.

### TRADITIOM longitudinal cell tracking

1×10^5^ differentially barcoded cells were expanded to reach ~90×10^6^ and plated based on the following scheme: 1) 31 hyperflasks were seeded (1.2×10^6^ and 2.8×10^6^ cells for -E2 and TAM conditions respectively), 2) 3×10^6^ cells were kept in culture as untreated arm (UT, 2 replicates), 3) 10×10^6^ were expanded to 90×10^6^ and harvested as triplicate POT samples, 4) 5×10^6^ cells were expanded to 40×10^6^ cells for plating of twenty time zero (T0) samples and collected after 48 hours, 5) The rest of the barcoded MCF7 population was frozen.

Cell treatment of the 31 aliases, with either -E2 or TAM, started 48 hours after seeding. Harvesting of each HYPERflask was performed at the indicated time points (shared timepoints for 2 replicates at day 7, day 14, 1 month, 2 months, 3 months and diverging time points for individual awakenings). At time of collection, cells were snap-frozen in multiple pellets for subsequent DNA and RNA extraction (for WGS, genomic-barcode sequencing and RNA-sequencing, respectively) or post-translational histone modification MS analysis. Following awakening, TAMα-**ζ** and AIα-ε samples were further cultured for 1 month in T150 flasks (Corning) with cell passaging giving rise to TEPs. Several aliquots of cells were frozen at awakening and TEP timepoints.

### Barcode amplification and next generation library preparation

Barcoded MCF7 cell lines were harvested and pelleted at indicated time points (POT, latency, dormancy, awakening and TEP). Genomic DNA isolation was performed using DNeasy Blood and Tissue DNA extraction kit (Qiagen) according to manufacturer’s recommendations. Qubit (Life Technologies) was used to quantify genomic DNA. Genomic barcode amplification was performed using Titanium Taq DNA polymerase (Clontech-Takare 639208) with a maximum of 50ng of DNA per reaction. When DNA extraction resulted in more than 50 ng, multiple reactions were performed to amplify the whole material and the PCR products were combined before library preparation. The following primer sequences were used for amplification:Fwd:ACCGAACGCAACGCACGCA, Rev:ACGACCACGACCGACCCGAACCACGA. TapeStation 2200 (Agilent) was used to detect 151-bp PCR amplicon including the 48-bp semi-random barcode sequence. After purification with SPRIselect beads (Beckman Coulter), NGS libraries were prepared using the NEBnext Ultra II DNA library preparation kit for Illumina (New England Biolabs) according to manufacturer’s recommendations. Libraries were detected and quantified using TapeStation and Qubit. NGS was performed at Novogene (Cambridge, UK) using NovaSeq6000 platform (Ilumina) (paired-end 150 bp).

### Drug response curves and proliferation assays

1000 cells per well from UT and TAM or -E2 TEPs were seeded in 96-well standard plates (Corning). Following overnight incubation, UT and TAM TEP cells were treated with 10-fold increasing concentrations of 4-OHT (1nM-10uM), vehicle control (EtOH) or re-exposure to E2 (10nM) in five independent replicates. On the other hand, UT and -E2 TEPs underwent several treatment conditions in five independent replicates: -E2, 100 nM TAM, 100nM Fulvestrant (Fulv, Sigma I4409), 50nM CDK7 inhibitor (CDK7i, kindly provided by Prof. Simak Ali), 100nM Palbociclib (Palbo, SIGMA PD 0332991) and re-exposure to E2 (10nM). Percentage of confluency was assessed and automatically calculated based on the images acquired with IncuCyte^®^ Zoom Live-Cell Analysis System (Sartorius) both at the day of compound addition (day 0) and at 7 days of incubation in a 37 °C and 5% CO_2_ cell culture incubator. Day 7 data were normalized to day 0 and overall data were represented as confluency fold changes over time.

### Genomic Barcodes Bioinformatic analysis

Raw reads were trimmed for adapters and quality (Phred Quality>=30) with trim_galore(version 0.6.4_dev)[cite with url https://www.bioinformatics.babraham.ac.uk/proiects/trim_galore/,author **<u>FelixKrueger</u>**]. After confirming the quality of processed reads with FastQC (version v0.11.9), they were mapped to the barcode reference library (10e6 barcode sequences) with BWA mem (version 0.7.17-r1188) [PMID: 19451168] using default settings. The alignment maps were then parsed with samtools (version 1.9) [PMID: 27536334] to filter out all the reads with supplementary alignments and alignment quality less than 30. The filtered and sorted alignment maps were used to count number of reads per barcode. These read counts were normalized with library size for each sample to get barcode frequencies. Survival dynamics were studied based on number of barcodes with non-zero frequency in each sample. Common barcodes in different pairs of awakening flasks/replicates were then checked to get percentages of overlapping barcodes in carbon copies from total surviving barcodes in each respective flask and also from the POT (number of barcodes existing in either of the 3 replicates). Barcodes were then tracked based on these frequency values to observe evolution dynamics with time and across different flasks. To visualize clonal evolution, fishplots were plotted using ‘stat_smooth’ function (geom= ‘area’, method = ‘loess’) of ggplot2 package in R (version 3.6.1) on the frequency values. Y-axis ranges from 0 to 1 value of frequency in both the directions, always adding up to 1.

We estimated doubling time for cells during the expansion phase of the experimental design, where we started from ~100,000 cells with 1 barcode per cell and get ~90 million cells with a broad range of frequency values after 13 days in culture under unperturbed conditions. Considering in ‘nt’ days a cell expands to 2^n-1^ cells, to reach ‘x’ number of cells in 13 days, the cells need ‘t’ days as described below:

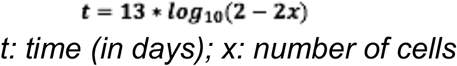

Doubling time in hours was calculated using this formula corresponding to all the barcodes in POT samples where relative number of cells were calculated with frequency values to add up to 90 million in each replicate.

### Whole Genome Sequencing (WGS) sample preparation and analysis

DNA was extracted using DNeasy Blood and Tissue DNA extraction kit (Qiagen) according to manufacturer’s recommendations. Qubit (Life Technologies) was used for quantification. Quality control and library preparation (28/30 samples prepared using PCR-free library protocol) were performed by Novogene (Cambridge, UK) where 150 bp paired-end sequencing (30X coverage) was performed on the Illumina NovaSeq6000 platform. Trim Galore [cite with url https://www.bioinformatics.babraham.ac.uk/projects/trim_galore/,author **FelixKrueger**] version 0.6.4 was used for adapter trimming of reads. Alignment to the hg38 human genome reference was performed using bwa-mem [PMID: 19451168] algorithm version 0.7.15. Conversion to binary, removal of PCR duplicates, sorting and indexing were performed using sambamba PMID: <u>25697820</u> version 0.7.0. Postprocessing and variant identification were performed using the Genome Analysis Toolkit PMID: 21478889, 20644199 (GATK) version 4.1.3.0 best practices: adding read groups using picard Wysoker A, Tibbetts K, Fennell T (2011). *icardTools* 1.5. 3. Available: http://sourceforge.net/projects/picard/files/picard-tools/. Accessed 2013 August 25 [<u>Google Scholar</u>] version 2.20.6, base quality recalibration using gatk BaseRecalibrator and gatk ApplyBQSR algorithms, and somatic variant calling using Mutect2 using as normal the POT sample pre-processed bam and the population germline resource af-only-gnomad.hg38.vcf.gz from the GATK resource bundle, with parameter –af-of-alleles-not-in-resource set as 0.001 and disabling MateOnSameContigOrNoMappedMateReadFilter filter. Variants were further filtered using gatk FilterMutectCalls and only PASS mutations were further processed.

### Noncoding analysis

Noncoding variants were selected via OpenCravat (all SNV excluding Exome). Noncoding SNVs were filtered with the ENCODE Cis Regulatory Element function and sorted for LINSIGHT score. Noncoding SNVs called in 3 carbon copies and with a LINSIGHT score >0.4 were overlapped with our unpublished noncoding CRISPR-KRAB Screen (Repression of CRE under oestrogen deprivation).

### RNA Sequencing analysis

Total RNA was extracted using QIAzol (Qiagen) and RNeasy Mini Kit (Qiagen). Quality control, mRNA library preparation (polyA enrichment) and sequencing were done at Novogene (Cambridge, UK) using NovaSeq6000 platform (paired-end 150 bp). Raw reads for all the RNA-Seq samples were trimmed for adapters and quality (Phred quality >= 30) with trim_galore (version 0.6.4_dev). After quality check with FastQC (version 0.11.5) the reads were pseudo-aligned to reference transcriptome (Homo_sapiens.GRCh38.96) with Kallisto (version 0.46.2) [PMID: 27043002]. Estimated transcript abundance values reported by ‘kallisto quant’ (number of bootstrap samples=100), were imported by ‘tximport’ package in R (version 3.6.1) and gene level summarization was performed using EnsDb.Hsapiens.v96 (ignoreAfterBar=T, ignoreTxVersion=T). DESeq dataset was then created with ‘DESeqDataSetFromTximport’ and lowly expressed genes were filtered based on having at least 10 counts in more than 3 samples. The filtered dataset was then normalized with variance stabilizing transformation (VST) to perform principal component analysis (PCA) using plotPCA function of DESeq2.

Differential expression for POT vs the samples at latent phase and Dormant vs the awakening samples was estimated with DESeq2 [PMID: 25516281]. Shrunken log fold changes were calculated, to identify differentially expressed genes (DEGs) in all logical comparisons, with the lfcShrink function using ‘apeglm’ as shrinkage estimator.

Genes with a fold change of at least 1.5x and adjusted p-values less than 0.01 were selected as significantly differentially expressed.

DESeq2 Wald statistic (stat) values were used to create the ranked lists genes based on the expression profiles. These were used to analyze gene set enrichments for up or down regulated genes in each comparison using GSEA software (version 3.0) [PMID: 16199517]. The enrichments were observed with a background of hallmark gene sets that represent well defined biological processes curated by aggregating various MSigDB gene sets (h.all.v7.4.symbols.gmt). Additionally, pre-adapted SWNE up and down signatures identified by Hong SP et al. were manually added to the hallmark gene sets before performing the enrichment analysis. Significantly enriched gene sets were reported with a false discovery rate (FDR) of 25 percent as threshold.

To compare awakening flasks and their TEPs (for which the replicates diverge a lot) we considered each flask independently. Missing the replicate information in this case, we estimated DEGs using edgeR package [PMID: 22287627] with recommended pipeline for samples without replicates. Common negative binomial dispersion was estimated with ‘estimateGLMCommonDisp’ function with robust option, ‘deviance’ method and without design model. Negative binomial generalized log-linear model was then fit to the read counts for each gene based on the defined contrast. This was followed by likelihood ratio test with ‘glmLRT’ function and the results were then parsed to keep DEGs with p-value of less than 0.001. Ranked gene list for GSEA in case of these comparisons was created using the reported p-value multiplied by sign of the fold change.

### Mass spectrometry (MS)

Sample preparation for histone post-translational modifications analysis by MS: Histones were enriched from 0.5-2×10^6^ cells as previously described (PMID: 31605746). The yield of histones was estimated by SDS-PAGE gel by comparison with known amounts of recombinant histone H3.1, following protein detection with colloidal Comassie staining. Approximately 2-4 μg of histones were mixed with a heavy-labelled histone super-SILAC mix, which was generated as previously described and used as an internal standard for quantification. (PMID: 28137569; PMID: 26463340). The samples were separated on an SDS-PAGE gel and in-gel digested. A band corresponding to the histone octamer (H3, H4, H2A, H2B) was excised, chemically acylated with propionic anhydride and in-gel digested with trypsin. After elution, the samples were derivatized with phenyl isocyanate (PMID: 25680960).

#### MS analysis of histone PTMs

Peptide mixtures were separated by reversed-phase chromatography on an EASY-Spray column (Thermo Fisher Scientic), 25-cm long (inner diameter 75 μm, PepMap C18, 2 μm particles), which was connected online to a Q Exactive Plus instrument through an EASY-Spray™ Ion Source (Thermo Fisher Scientific). Solvent A was 0.1% formic acid (FA) in ddH2O and solvent B was 80% ACN plus 0.1% FA. Peptides were injected in an aqueous 1% TFA solution at a flow rate of 500 nl/min and were separated with a 50-min linear gradient of 10–45%. The Q Exactive instrument was operated in the data-dependent acquisition (DDA) mode. Survey full scan MS spectra (m/z 300–1350) were analyzed in the Orbitrap detector with a resolution of 60,000-70,000 at m/z 200. The 12 most intense peptide ions with charge states comprised between 2 and 4 were sequentially isolated to a target value for MS1 of 3×10^6^ and fragmented by HCD with a normalized collision energy setting of 28%. The maximum allowed ion accumulation times were 20 ms for full scans and 80 ms for MS/MS, and the target value for MS/MS was set to 1×10^5^. The dynamic exclusion time was set to 10 s, and the standard mass spectrometric conditions for all experiments were as follows: spray voltage of 1.8 kV, no sheath and auxiliary gas flow. The acquired RAW data were analyzed using Epiprofile 2.0 (PMID: 29790754), selecting the SILAC option, followed by manual validation (PMID: 29790754). For each histone modified peptide, the % relative abundance (%RA) was estimated by dividing the area under the curve (AUC) of each modified peptide for the sum of the areas corresponding to all the observed forms of that peptide and multiplying by 100. Light/Heavy (L/H) ratios of %RA were then calculated. Data display was carried out using Perseus (PMID: 27348712). Normalized L/H ratios, defined as L/H ratios of relative abundances normalized over the average value across the samples, were visualized and clustered with correlation distance and average linkage as parameters.

## Acknowledgements

We are immensely grateful to all the patients and their families for donating clinical samples for the study and for supporting cancer research. We want to acknowledge Charing Cross (CX) Hospital (London, UK), the IRST (Meldola, IT) and all the personnel involved in the day-to-day clinical work. This work is funded by grants from CRUK (A23110, L.M, D.R and consumables) and by the Katherine and Douglas Longden legacy (E.S, H.D and consumables) and infrastructure support was provided by Imperial Experimental Cancer Medicine Centre, Cancer Research UK Imperial Centre, National Institute for Health Research (NIHR) Imperial Biomedical Research Centre (BRC) and Imperial College Healthcare NHS Trust Tissue Bank. We like to acknowledge Iros Barozzi, Phillipp Thomas, Andrea Sottoriva, Massimiliano Bonafe and the METAMORPHEUS team-members for all the stimulating and critical discussion. The answer to dormancy is 42.

## Supplementary information

**Supplementary figure 1:**
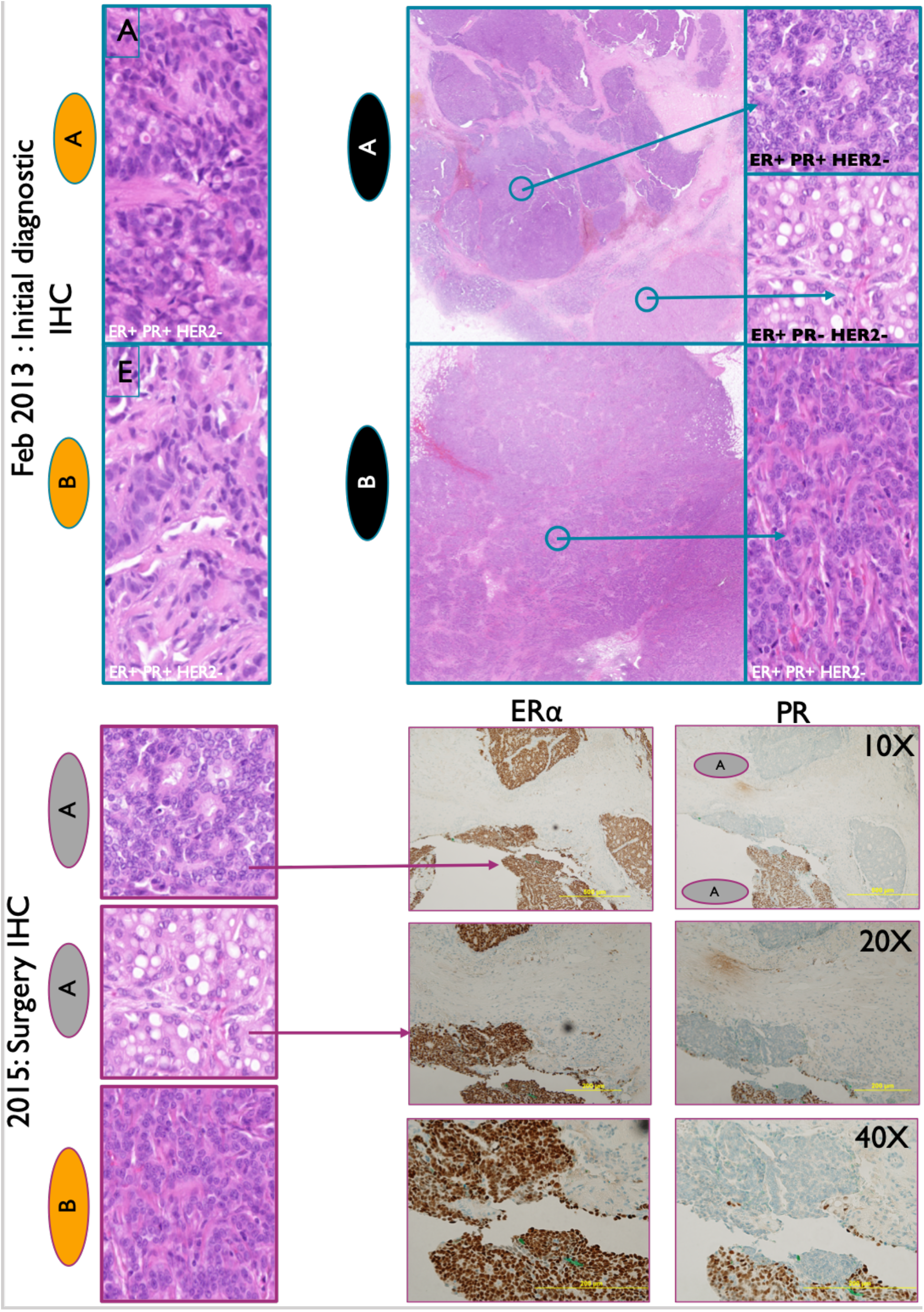
Representative images of H&E and immunohistochemical (IHC) staining of tumors from Patients A and B at diagnostic (upper panel) and surgical biopsies (lower panel).

**Supplementary figure 2:**
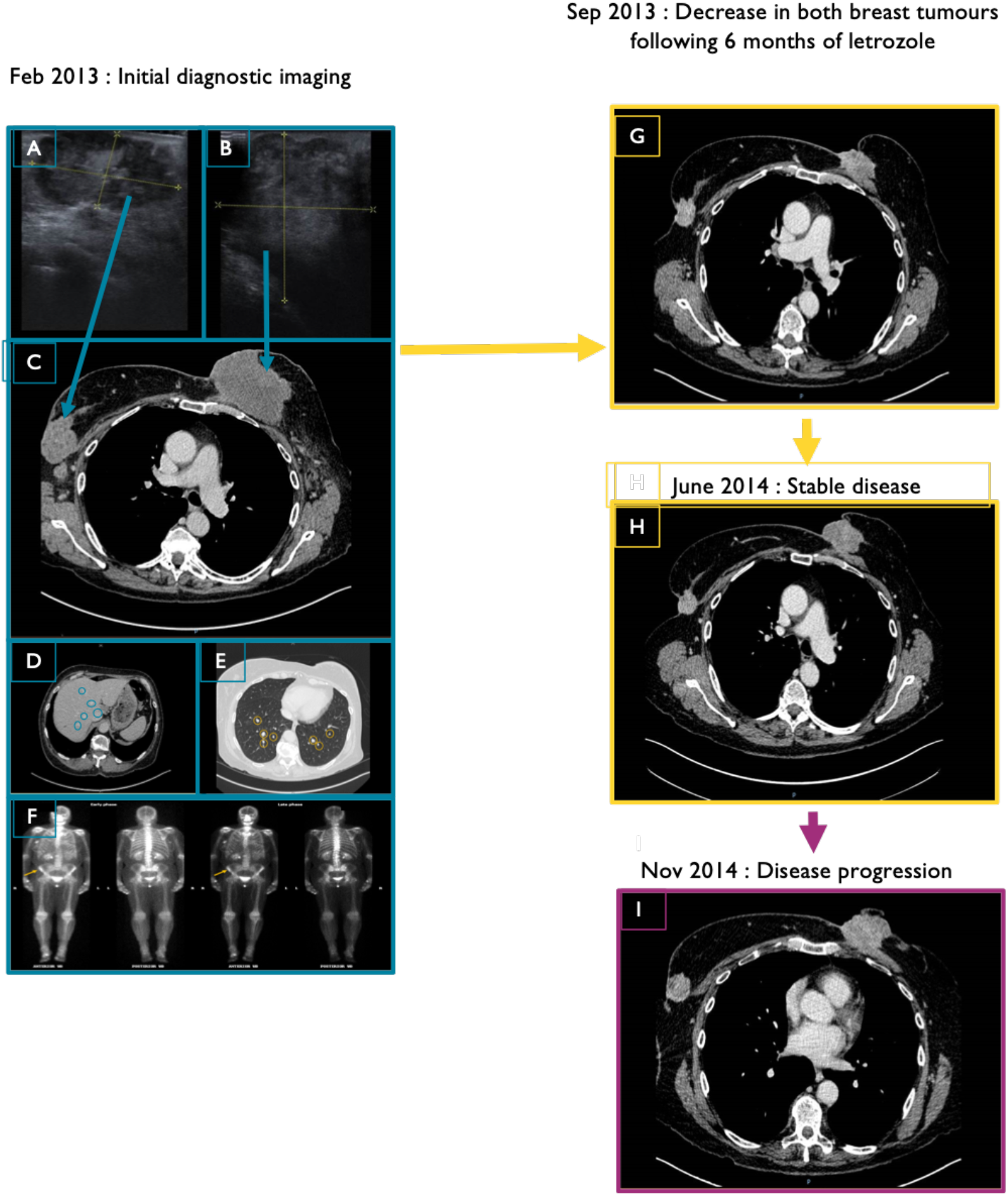
Ultrasound scan of patient A at diagnosis and after 6 months of ET (letrozole). **A.** (February 2013) Ultrasound scan showing the right breast lesion measuring 50.9 mm in maximal diameter; **B.** Ultrasound scan showing the fungating left breast lesion measuring 62.7mm x 58.5mm; **C.** Axial section of an intravenous (I.V.) contrast phase computer tomography (CT) scan of the thorax revealing right (52 mm x 41 mm) and left (82 mm x 63 mm) breast tumours; **D.** Axial section of an I.V. contrast phase CT scan of the liver showing liver metastases highlighted by the yellow circles; **E.** Axial section of an I.V. contrast phase CT scan of the thorax showing bilateral lung nodules suggesting lung metastases; **F.** A Nuclear medicine whole body bone scan showing uptake of radioactive tracer in the right iliac crest on early and late phase imaging indicative of a solitary sclerotic bone metastatic lesion. The patient was commenced on neoadjuvant letrozole and a 6 month repeat CT scan (**G)** revealed disease response with the left tumour now measuring 49 mm x 24 mm and the right tumour 30 mm x 24 mm (September 2013). A follow-up CT scan in June 2014 (**H)** showed stable disease with minimal shrinkage of both left and right breast tumours. In December 2014a further I.V. contrast CT scan. **(I)** revealed evidence of disease progression with the left tumour now measuring 62 mm and the right 33 mm in maximal diameter. The patient was then placed on the RADICAL trial with the addition of an FGFR inhibitor. She was then scheduled for a mastectomy and wide local excision for the left and right breast tumours, respectively.

**Supplementary figure 3:**
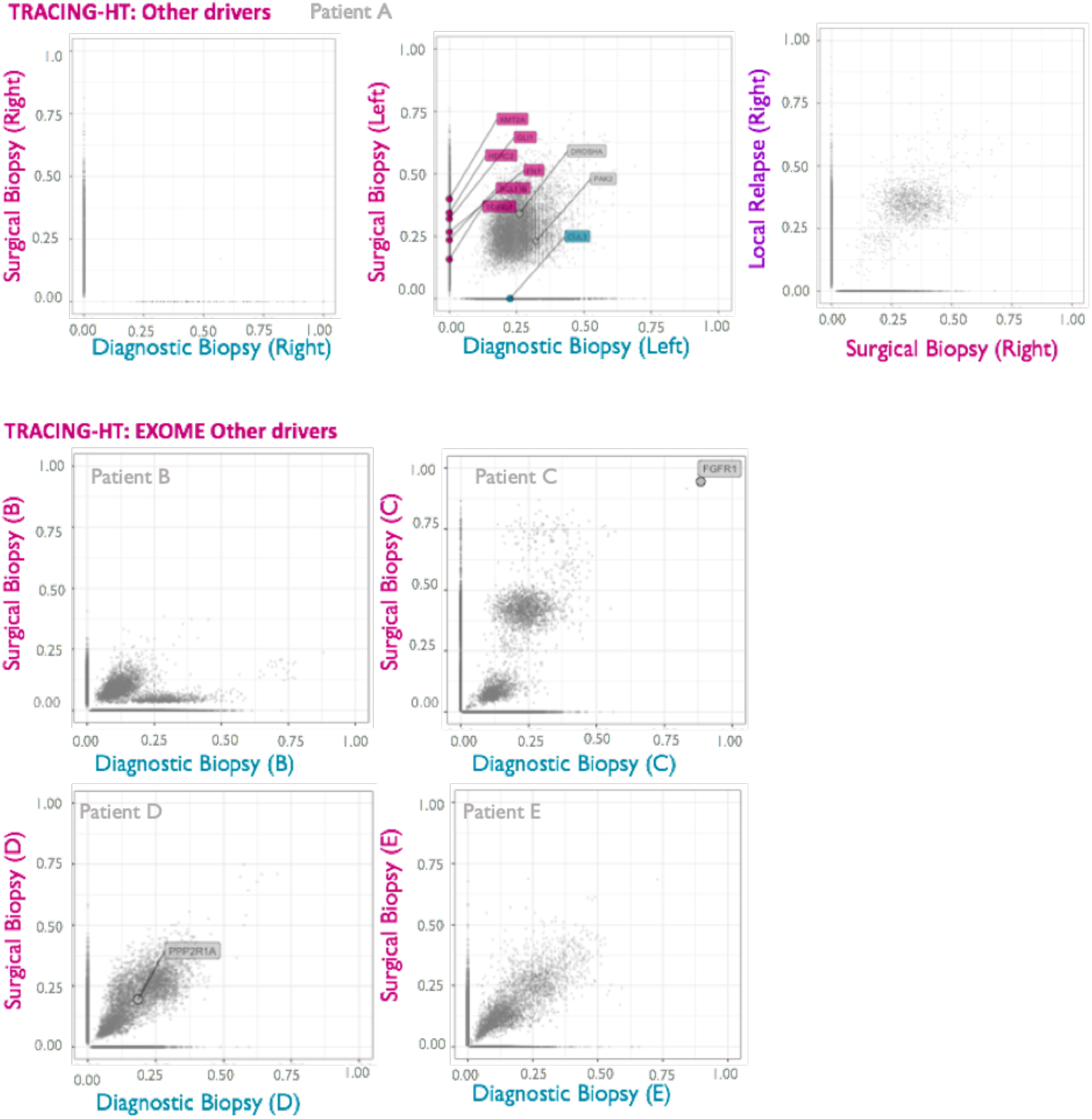
Variant Allele Frequency plots for all known cancer drivers other than breast cancer. The analysis was performed applying less stringent mutation filters (all cancer drivers and FATHMM significant score >0.6). All the variants identified within a sample are depicted by a grey point corresponding to their frequency value. Variants detected in driver genes are labeled and highlighted according to detection in surgical biopsy (pink), diagnostic biopsy (cyan) or both (grey).

**Supplementary figure 4:**
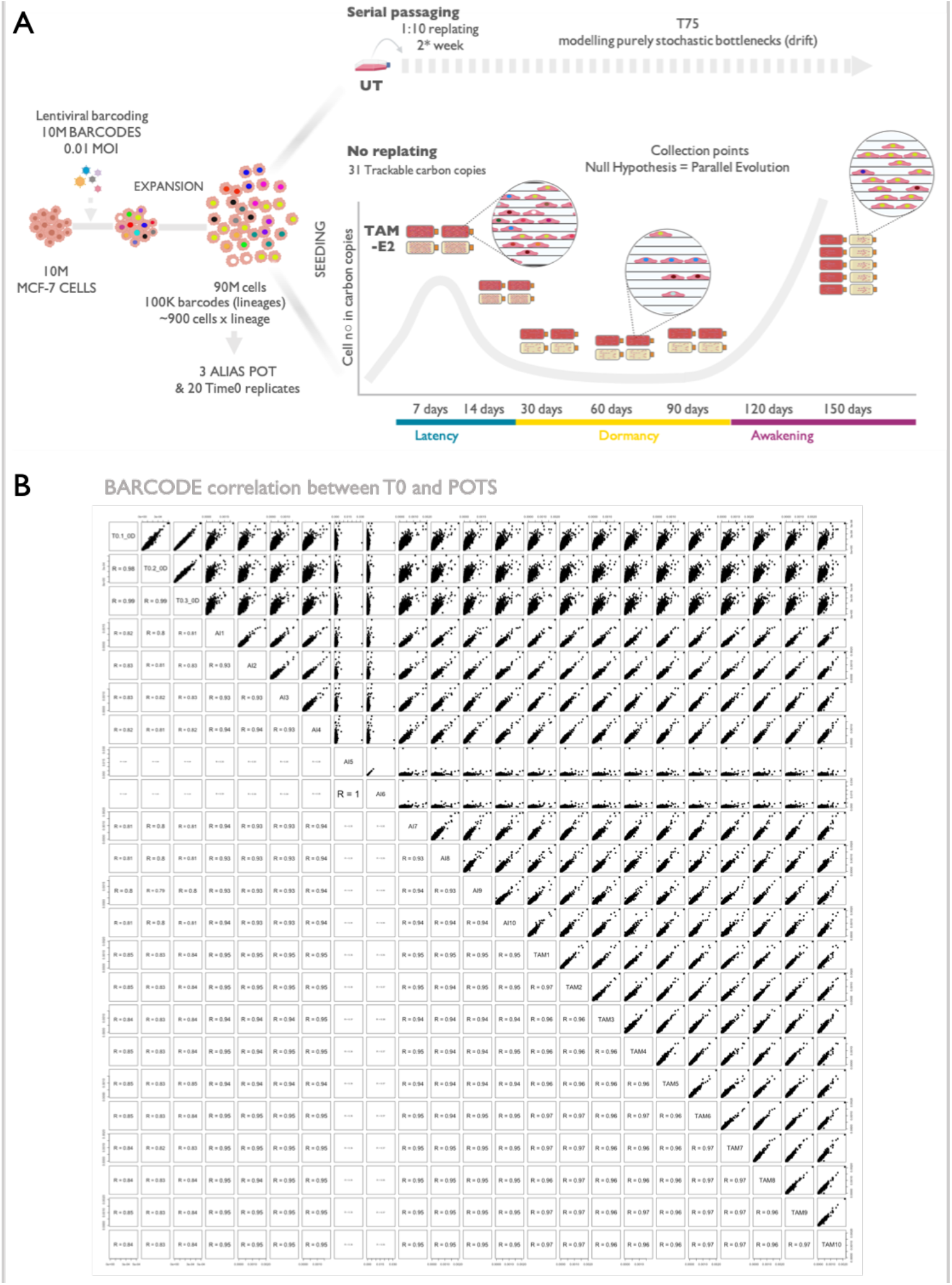
Experimental design. **A.** Ten million MCF7 breast cancer cells were lentivirally barcoded with a plasmidic library containing ten million individual barcodes. Using a 0.01 MOI we obtained a founder population of 1*10^5^ individually barcoded cells. Cells were selected and expanded to a population of 90 million cells (POT). 31 replicates were seeded in HYPERflasks with ~2 million cells each and treated after 48 hours with either TAM or AI. Cells were never split along the whole experimental course and harvesting of each HYPERflask was performed at the indicated time points (shared time points for 2 replicates for latency and dormancy and diverging time points for individual awakenings). Remaining POT cells were either kept in culture as UT counterpart (and collected at the indicated timepoints), collected as POT triplicate or seeded as 20 time zero (T0) replicates and collected after 48 hours. **B.** Correlation analysis between barcode composition in the 3 POTs and in the 20 time zero (T0) replicates.

**Supplementary figure 5:**
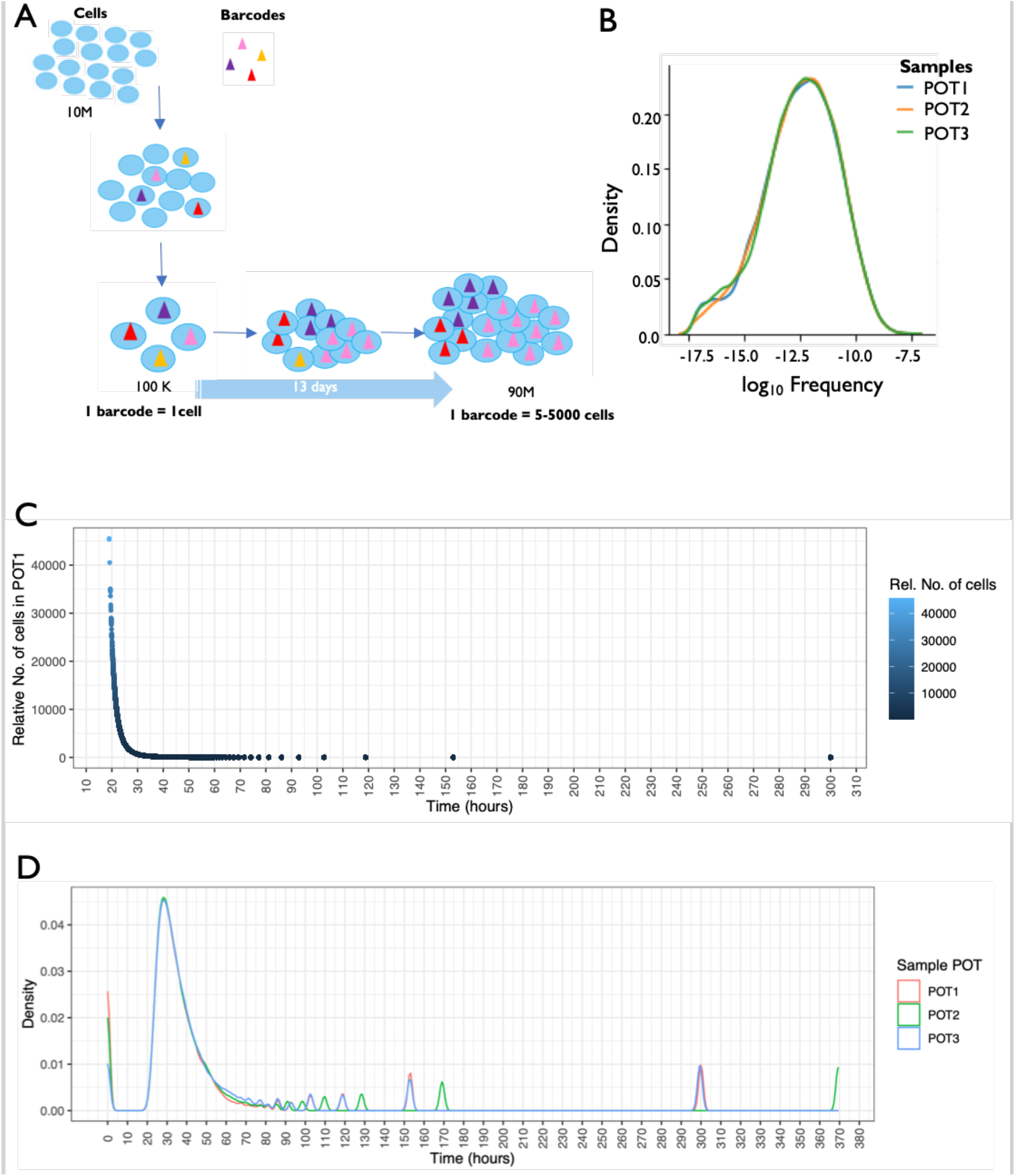
Mathematical modelling representing heterogeneous cell cycle dynamics. **A.** A pictorial representation showing the barcoding process followed by selection and expansion for 13 days. Ultra-low MOI of 0.01 ensures 1 barcode per cell but during expansion there is no uniformity in the growth rates resulting in wide range of frequency per barcode. **B.** Barcode frequency distribution in 3 POT replicates confirms a big range of frequency values in 13 days despite starting from equally represented cells. **C.** Relative number of cells in POT1 plotted across estimated doubling time in hours indicate presence of very slow cycling cells that might have undergone cell cycle only twice in 13 days. **D.** Density plot of relative cell abundance across estimated doubling time shows consistency in 3 POT replicates.

**Supplementary figure 6:**
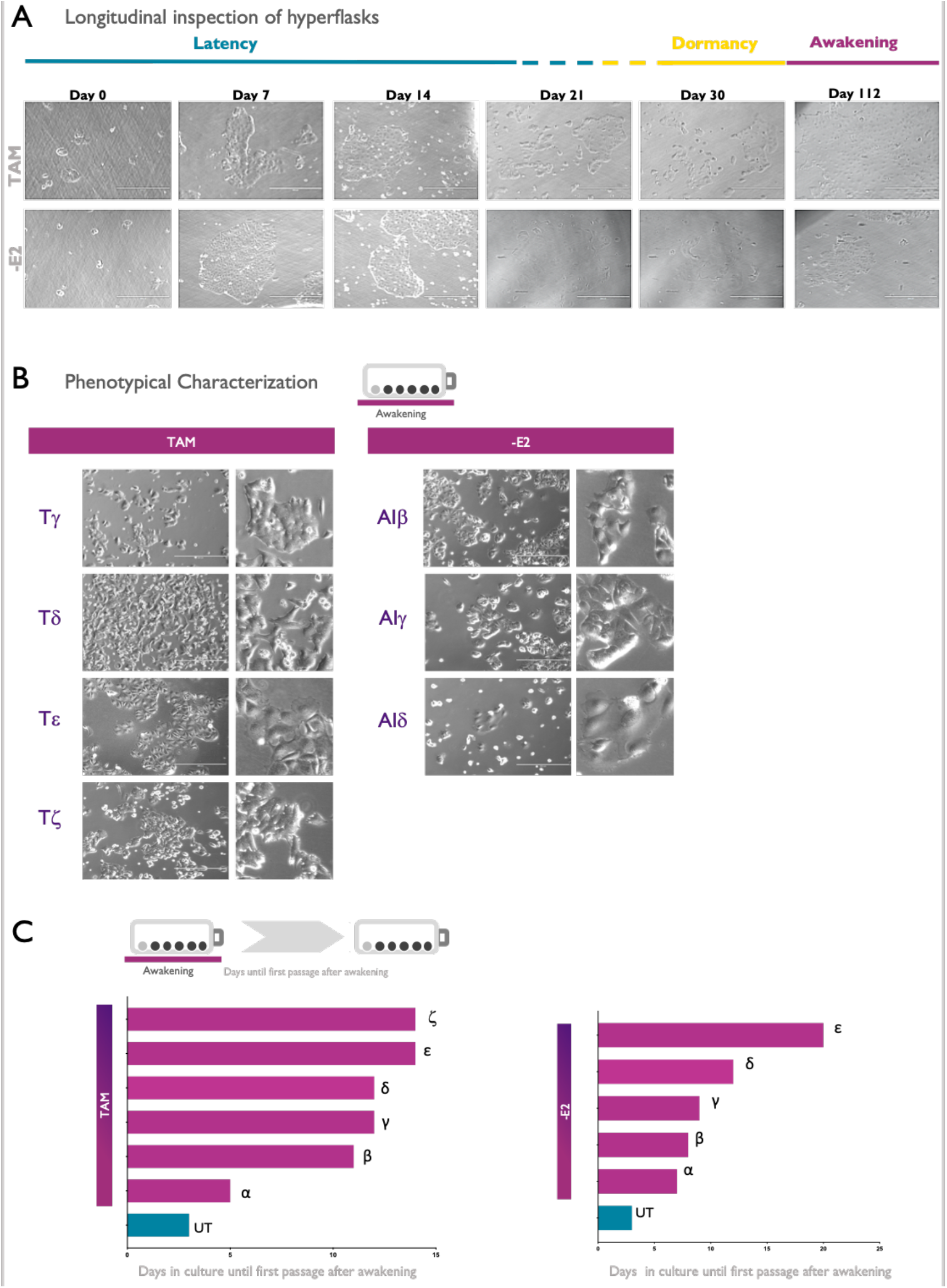
Morphological inspection of cells along adaptation. **A.** Representative bright-field images of –E2 and TAM hyperflasks (observable layers) along latency, dormancy and awakening. Images were captured with EVOS Cell Imaging Systems (10X). **B.** Morphological differences between untreated (UT) and awakening TEPs were captured with EVOS Cell Imaging Systems (10X). **C.** Number of days before the first passage of awakening cells detached from hyperflasks and replated in standard culture conditions.

**Supplementary figure 7:**
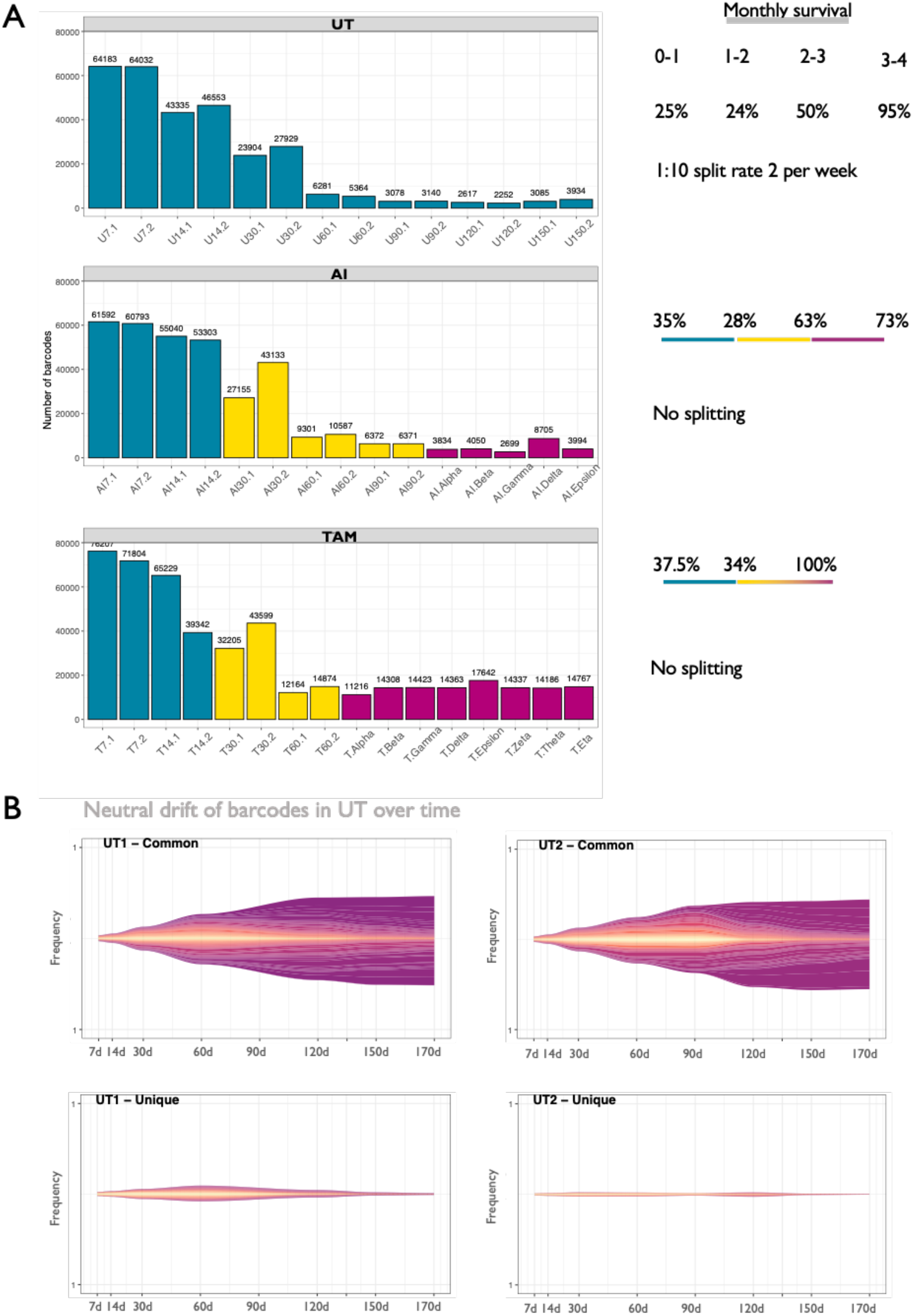
Barcode extinction dynamics. **A.** Barcode counts, at each indicated time point, are listed for UT, AI (-E2) and TAM arms. A summary of the percentages of surviving barcodes along the collection points is shown on the right of the panel. **B.** Frequencies of common and unique barcodes in the UT condition along the collection points.

**Supplementary figure 8:**
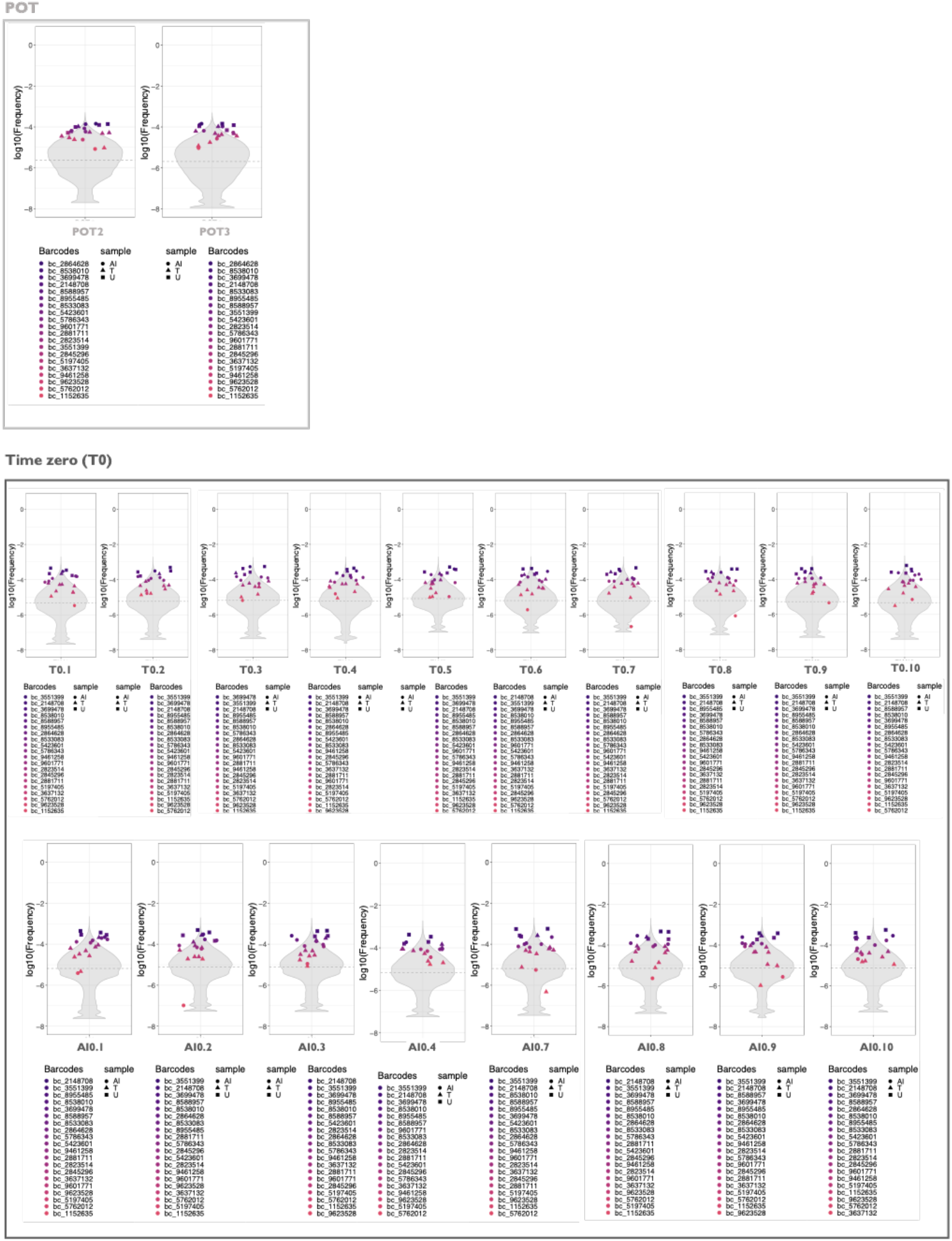
Violin plots depicting the frequency distribution of barcodes in POT and time zero (T0) carbon copies. Winning barcodes among all the treatment conditions (Fig. 4A) are indicated under each violin plot in order of frequency. Specific symbols (square for UT, circle for -E2 and triangle for Tam) indicate the belonging of the winning barcodes to the respective treatment condition.

**Supplementary figure 9:**
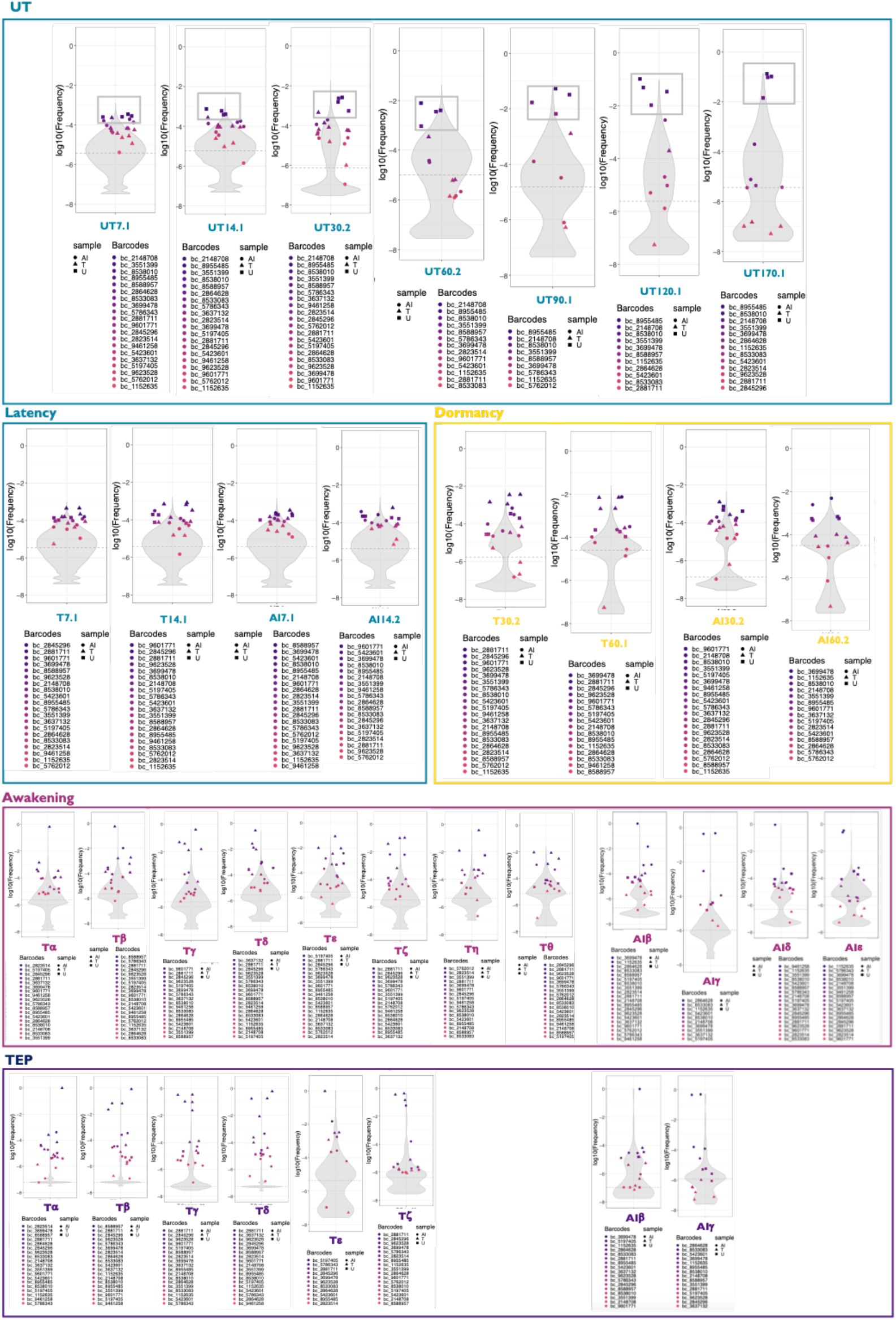
Violin plots depicting the frequency distribution of barcodes in carbon copies, during their adaptive journey through latency, dormancy, awakening and TEPs for –E2 and TAM conditions along with UT counterpart. Winning barcodes among all the treatment conditions (Fig. 4A) are indicated under each violin plot in order of frequency at that specific collection point. Specific symbols (square for UT, circle for -E2 and triangle for Tam) indicate the belonging of the winning barcodes to the respective treatment condition.

## References

Acar, A., Nichol, D., Fernandez-Mateos, J., Cresswell, G.D., Barozzi, I., Hong, S.P., Trahearn, N., Spiteri, I., Stubbs, M., Burke, R., et al. (2020). Exploiting evolutionary steering to induce collateral drug sensitivity in cancer. Nat Commun 11, 1923.

Ben-David, U., Siranosian, B., Ha, G., Tang, H., Oren, Y., Hinohara, K., Strathdee, C.A., Dempster, J., Lyons, N.J., Burns, R., et al. (2018). Genetic and transcriptional evolution alters cancer cell line drug response. Nature 560, 325–330.

Bertucci, F., Ng, C.K.Y., Patsouris, A., Droin, N., Piscuoglio, S., Carbuccia, N., Soria, J.C., Dien, A.T., Adnani, Y., Kamal, M., et al. (2019). Genomic characterization of metastatic breast cancers. Nature 569, 560–564.

Chen, F., Ding, K., Priedigkeit, N., Elangovan, A., Levine, K.M., Carleton, N., Savariau, L., Atkinson, J.M., Oesterreich, S., and Lee, A.V. (2021). Single-Cell Transcriptomic Heterogeneity in Invasive Ductal and Lobular Breast Cancer Cells. Cancer Res 81, 268–281.

Chung, J.H., Pavlick, D., Hartmaier, R., Schrock, A.B., Young, L., Forcier, B., Ye, P., Levin, M.K., Goldberg, M., Burris, H., et al. (2017). Hybrid capture-based genomic profiling of circulating tumor DNA from patients with estrogen receptor-positive metastatic breast cancer. Ann Oncol 28, 2866–2873.

Hinohara, K., Wu, H.-J., Vigneau, S., McDonald, T.O., Igarashi, K.J., Yamamoto, K.N., Madsen, T., Fassl, A., Egri, S.B., Papanastasiou, M., et al. (2018). KDM5 Histone Demethylase Activity Links Cellular Transcriptomic Heterogeneity to Therapeutic Resistance. Cancer Cell.

Hong, S.P., Chan, T.E., Lombardo, Y., Corleone, G., Rotmensz, N., Bravaccini, S., Rocca, A., Pruneri, G., McEwen, K.R., Coombes, R.C., et al. (2019). Single-cell transcriptomics reveals multi-step adaptations to endocrine therapy. Nat Commun 10, 3840.

Horvath, S. (2013). DNA methylation age of human tissues and cell types. Genome Biology 14, 3156.

Liu, M., Ohtani, H., Zhou, W., Ørskov, A.D., Charlet, J., Zhang, Y.W., Shen, H., Baylin, S.B., Liang, G., Grønbæk, K., et al. (2016). Vitamin C increases viral mimicry induced by 5-aza-2’-deoxycytidine. Proc National Acad Sci 113, 10238–10244.

Magnani, L., Eeckhoute, J., and Lupien, M. (n.d.). Pioneer factors: directing transcriptional regulators within the chromatin environment. 27.

Magnani, L., Stoeck, A., Zhang, X., Lánczky, A., Mirabella, A.C., Wang, T.-L., Gyorffy, B., and Lupien, M. (2013). Genome-wide reprogramming of the chromatin landscape underlies endocrine therapy resistance in breast cancer. Proc National Acad Sci 110, E1490–E1499.

Magnani, L., Frigè, G., Gadaleta, R., Corleone, G., Fabris, S., Kempe, H., Verschure, P.J., Barozzi, I., Vircillo, V., Hong, S.-P., et al. (2017). Acquired CYP19A1 amplification is an early specific mechanism of aromatase inhibitor resistance in ERα metastatic breast cancer. Nature Genetics 49, 444.

Martincorena, I., Fowler, J.C., Wabik, A., Lawson, A.R., Abascal, F., Hall, M.W., Cagan, A., Murai, K., Mahbubani, K., Stratton, M.R., et al. (2018). Somatic mutant clones colonize the human esophagus with age. Science 362, eaau3879.

Martínez-Jiménez, F., Muiños, F., Sentís, I., Deu-Pons, J., Reyes-Salazar, I., Arnedo-Pac, C., Mularoni, L., Pich, O., Bonet, J., Kranas, H., et al. (2020). A compendium of mutational cancer driver genes. Nat Rev Cancer 20, 555–572.

Nguyen, V.T.M., Barozzi, I., Faronato, M., Lombardo, Y., Steel, J.H., Patel, N., Darbre, P., Castellano, L., Győrffy, B., Woodley, L., et al. (2015). Differential epigenetic reprogramming in response to specific endocrine therapies promotes cholesterol biosynthesis and cellular invasion. Nat Commun 6, 10044.

Nik-Zainal, S., Alexandrov, L.B., Wedge, D.C., Loo, P., Greenman, C.D., Raine, K., Jones, D., Hinton, J., Marshall, J., Stebbings, L.A., et al. (2012). Mutational Processes Molding the Genomes of 21 Breast Cancers. Cell 149, 979–993.

Nik-Zainal, S., Davies, H., Staaf, J., Ramakrishna, M., Glodzik, D., Zou, X., Martincorena, I., Alexandrov, L.B., Martin, S., Wedge, D.C., et al. (2016). Landscape of somatic mutations in 560 breast cancer whole-genome sequences. Nature 534, 47.

Pan, H., Gray, R., Braybrooke, J., Davies, C., Taylor, C., McGale, P., Peto, R., Pritchard, K.I., Bergh, J., Dowsett, M., et al. (2017). 20-Year Risks of Breast-Cancer Recurrence after Stopping Endocrine Therapy at 5 Years. New Engl J Medicine 377, 1836–1846.

Petljak, M., Alexandrov, L.B., Brammeld, J.S., Price, S., Wedge, D.C., Grossmann, S., Dawson, K.J., Ju, Y.S., Iorio, F., Tubio, J.M.C., et al. (2019). Characterizing Mutational Signatures in Human Cancer Cell Lines Reveals Episodic APOBEC Mutagenesis. Cell 176, 1282–1294.e20.

Phan, T.G., and Croucher, P.I. (2020). The dormant cancer cell life cycle. Nat Rev Cancer 1–14.

Razavi, P., Chang, M.T., Xu, G., Bandlamudi, C., Ross, D.S., Vasan, N., Cai, Y., Bielski, C.M., Donoghue, M., Jonsson, P., et al. (2018). The Genomic Landscape of Endocrine-Resistant Advanced Breast Cancers. Cancer Cell 34, 427–438.e6.

Rehman, S.K., Haynes, J., Collignon, E., Brown, K.R., Wang, Y., Nixon, A.M.L., Bruce, J.P., Wintersinger, J.A., Mer, A.S., Lo, E.B.L., et al. (2021). Colorectal Cancer Cells Enter a Diapause-like DTP State to Survive Chemotherapy. Cell 184, 226–242.e21.

Roulois, D., Loo Yau, H., Singhania, R., Wang, Y., Danesh, A., Shen, S.Y., Han, H., Liang, G., Jones, P.A., Pugh, T.J., et al. (2015). DNA-Demethylating Agents Target Colorectal Cancer Cells by Inducing Viral Mimicry by Endogenous Transcripts. Cell 162, 961–973.

Sansone, P., Ceccarelli, C., Berishaj, M., Chang, Q., Rajasekhar, V.K., Perna, F., Bowman, R.L., Vidone, M., Daly, L., Nnoli, J., et al. (2016). Self-renewal of CD133hi cells by IL6/Notch3 signalling regulates endocrine resistance in metastatic breast cancer. Nat Commun 7, 10442.

Shihab, H.A., Rogers, M.F., Gough, J., Mort, M., Cooper, D.N., Day, I.N.M., Gaunt, T.R., and Campbell, C. (2015). An integrative approach to predicting the functional effects of non-coding and coding sequence variation. Bioinformatics 31, 1536–1543.

Siersbæk, R., Scabia, V., Nagarajan, S., Chernukhin, I., Papachristou, E.K., Broome, R., Johnston, S.J., Joosten, S.E.P., Green, A.R., Kumar, S., et al. (2020). IL6/STAT3 Signaling Hijacks Estrogen Receptor α Enhancers to Drive Breast Cancer Metastasis. Cancer Cell.

Suva, M.L., Riggi, N., and Bernstein, B.E. (2013). Epigenetic Reprogramming in Cancer. Science 339, 1567–1570.

Tomasetti, C., and Vogelstein, B. (2015). Variation in cancer risk among tissues can be explained by the number of stem cell divisions. Science 347, 78–81.

Tomasetti, C., Vogelstein, B., and Parmigiani, G. (2013). Half or more of the somatic mutations in cancers of self-renewing tissues originate prior to tumor initiation. Proc National Acad Sci 110, 1999–2004.

Tomasetti, C., Li, L., and Vogelstein, B. (2017). Stem cell divisions, somatic mutations, cancer etiology, and cancer prevention. Science 355, 1330–1334.

Toy, W., Carlson, K., Martin, T., Razavi, P., Berger, M., Baselga, J., Greene, G., Katzenellenbogen, J., and Chandarlapaty, S. (2019). Abstract P5-04-11: Non-canonical, clinical ESR1 mutations promote resistance to antiestrogen therapies. Poster Sess Abstr P5-04-11-P5-04-11.

Zhang, W., Bado, I.L., Hu, J., Wan, Y.-W., Wu, L., Wang, H., Gao, Y., Jeong, H.-H., Xu, Z., Hao, X., et al. (2021). The bone microenvironment invigorates metastatic seeds for further dissemination. Cell.

